# Genome-Wide Reconstitution of Chromatin Transactions: RSC Preferentially disrupts H2A.Z-Containing Nucleosomes

**DOI:** 10.1101/394692

**Authors:** Aylin Cakiroglu, Cedric R. Clapier, Andreas H. Ehrensberger, Elodie Darbo, Bradley R. Cairns, Nicholas M. Luscombe, Jesper Q. Svejstrup

**Affiliations:** Bioinformatics and Computational Biology Laboratory, 1 Midland Road, London, NW1 1AT, UK.; Department of Oncological Sciences, Huntsman Cancer Institute, and Howard Hughes Medical Institute, University of Utah School of Medicine, Salt Lake City, UT 84112, USA.; Mechanisms of Transcription Laboratory, The Francis Crick Institute, 1 Midland Road, London, NW1 1AT, UK.; UCL Genetics Institute, Department of Genetics, Evolution and Environment, University College London, Gower Street, London WC1E 6BT, UK.

## Abstract

Chromatin transactions are typically studied *in vivo*, or *in vitro* using artificial chromatin lacking the epigenetic complexity of the natural material. Attempting to bridge the gap between these approaches, we established a system for isolating the yeast genome as a library of mono-nucleosomes harboring the natural epigenetic signature, suitable for biochemical manipulation. Combined with deep sequencing, this library was used to investigate the intrinsic stability of individual nucleosomes, and – as proof of principle - the nucleosome preference of the chromatin remodeling complex, RSC. Our data indicate that the natural stability of nucleosomes differs greatly, with nucleosomes on tRNA genes and on promoters of protein-coding genes standing out as intrinsically unstable. Interestingly, RSC shows a distinct preference for nucleosomes derived from regions with a high density of histone variant H2A.Z, and this preference is indeed markedly diminished using nucleosomes from cells lacking H2A.Z. Importantly, the preference for H2A.Z remodeling/nucleosome ejection can also be reconstituted with recombinant nucleosome arrays. Together, our data indicate that, despite being separated from their genomic context, individual nucleosomes can retain their original identity as promoter- or TSS-nucleosomes. Besides shedding new light on nucleosome stability and the chromatin remodeler RSC, the simple experimental system outlined here should be generally applicable to the study of chromatin transactions.

## INTRODUCTION

The eukaryotic genome is stored as a polymer of nucleosomes in which ∼147 base pairs (bp) of DNA is wrapped around a core of eight histone proteins separated by a linker of 5-35 bp (Kornberg 1974; Kornberg and Lorch 1999). Because these complexes encompass the entire genome, they serve as the substrate and platform for all nuclear processes involving DNA. Nucleosomes influence reactions on DNA in at least two ways: by occluding access to the DNA they can exert an inhibitory function, and by recruiting enzymes and regulatory factors they may stimulate catalysis. The regulatory capacity of nucleosomes is amplified vastly through regional incorporation of an extensive assortment of post-translational histone marks and histone variants, in turn rendering every nucleosome unique (Strahl and Allis 2000; Turner 2000).

A long-standing challenge in the field of chromatin research has been the lack of a reconstituted genome-wide system for studying the role of epigenetic modifications in the context of their natural sequences. Current approaches typically involve either assembling chromatin *in vitro* from naked DNA and free histones, or studying it inside the cell, *in vivo*. Even though both approaches have substantially advanced our understanding of chromatin function in general, they also suffer some limitations: reconstituted chromatin has virtually none of the complexity of natural chromatin, being devoid of the natural genomic sequences and/or the combinatorial assortment of histone marks. Conversely, chromatin studied *in vivo* obviously retains its natural composition, but is trapped in a context in which a process of interest is continuously subjected to the direct or indirect influence of the numerous biochemical reactions taking place inside the cell. Protocols in which chromatin is isolated directly from the host organism in a manner suitable for biochemical reconstitution have been described, but they are generally limited to the study of one or a few loci at a time (Griesenbeck et al. 2003; Unnikrishnan et al. 2012; Hamperl et al. 2014; Ehrensberger et al. 2015). As such, a system in which an entire genome is available in native form for the biochemical reconstitution of chromatin transactions would be very useful.

RSC (Remodels the Structure of Chromatin) is a member of the Swi2/Snf2 family of ATP-dependent chromatin remodelers and the most abundant chromatin remodeling factor in yeast (Cairns et al. 1996). It is necessary for transcription by all three nuclear RNA polymerases (Parnell et al. 2008) and contributes to the establishment of the canonical nucleosome-depleted or ‘nucleosome-free’ region (NFR) found in the majority of yeast RNAPII promoters (Hartley and Madhani 2009; Wippo et al. 2011; Lorch et al. 2014). By itself, RSC destabilizes nucleosomes such that their DNA becomes sensitive to nuclease digestion (Cairns et al. 1996). In the presence of a histone acceptor such as naked DNA or the histone chaperone Nap1, RSC can fully disassemble mono-nucleosomes into naked DNA (Lorch et al. 1999; Lorch et al. 2006), but it can also eject nucleosomes from nucleosome arrays even in the absence of NAP1 (Clapier et al. 2016). The mechanism of RSC action has been studied through a variety of biochemical and structural approaches and involves RSC conducting directional DNA translocation from a site within the nucleosome, pumping DNA around the octamer resulting in nucleosome sliding and/or ejection of either the RSC-bound octamer, or the one adjacent to it (Saha et al. 2005; Chaban et al. 2008). Despite the abundance of data on its mechanism, the mode by which RSC is targeted to specific loci remains unclear. It can be recruited by transcription factors (Swanson et al. 2003; Inai et al. 2007), but it also harbors its own DNA-binding domains (Angus-Hill et al. 2001; Badis et al. 2008) and eight bromo-domains, at least one of which has been implicated in recruitment to acetylated histones (Kasten et al. 2004). RSC has been shown to selectively remodel at the promoter, but not the open-reading frame nucleosomes on purified *PHO5* gene rings (Lorch et al. 2011). It also shows a preference for nucleosomes bearing poly(dAdT) tracts *in vitro* (Lorch et al. 2014). We chose to work with RSC as the model enzyme to validate the functionality of genomic nucleosomes for several reasons: (1) when combined with NAP1, the reaction product, naked DNA, can easily be separated from mono-nucleosomes that are not remodeled for deep sequencing in order to characterize the underlying DNA, (2) the enzyme is abundant and the reaction robust, and (3) RSC plays important roles in transcription (Parnell et al. 2008; Parnell et al. 2015). The purpose of our approach was to identify nucleosomes that were disassembled preferentially above the generic background of RSC activity in the hope that we would gain new insights into the catalytic preferences of this important chromatin remodeler.

Histone H2A.Z (encoded by the non-essential *HTZ1* gene in budding yeast) is a variant of histone H2A that shares 60% sequence identity with its canonical H2A counterpart (Suto et al. 2000). While H2A.Z localizes almost exclusively to promoters (Zhang et al. 2005), and often occupies the two nucleosomes surrounding the NFR (Raisner et al. 2005), yeast cells lacking H2A.Z still contain intact NFRs (Hartley and Madhani 2009). Even though H2A.Z localizes to virtually all promoters, irrespective of expression level (Raisner et al. 2005), its occupancy differs between promoters (Albert et al. 2007). When genes are dynamically activated, H2A.Z occupancy decreases, suggesting a poising function that might assist in gene activation (Zhang et al. 2005). Incorporation of H2A.Z is catalyzed by the chromatin remodeler SWR1 by replacement of the H2A/H2B dimer on a canonical nucleosome with a H2A.Z/H2B dimer (Mizuguchi et al. 2004). Despite the remarkably distinctive localization of H2A.Z and the comprehensive understanding of its deposition mechanism, relatively little is known about the function of H2A.Z on the promoters on which it resides.

Nucleosome occupancies are determined not only by chromatin remodeling enzymes, but also by the intrinsic physical stability of nucleosomes. Features such as high AT-content and histone acetylation have thus been correlated with decreased nucleosome stability *in vitro* (Li et al. 1993; Brower-Toland et al. 2005; Tillo and Hughes 2009) and decreased occupancy *in vivo* (Yuan et al. 2005; Kaplan et al. 2009).

Here we show that a library of purified, native mono-nucleosomes isolated from yeast can be used to study the specificity of chromatin-modifying enzymes such as RSC, as well as the structural features of individual nucleosomes. Among other findings, this system has uncovered intrinsically highly unstable nucleosomes on tRNA genes, and a preference of RSC for ejection of H2A.Z-containing nucleosomes.

## Results

### Purification of genomic chromatin and description of assays

In order to generate a library of native yeast nucleosomes, we developed a three-step purification protocol (Figure 1A): first, purified yeast nuclei were incubated with micrococcal nuclease (MNase), which preferentially digests naked DNA to generate short chromatin fragments. The resulting fragments were extracted from the nuclei, then bound to and eluted from DEAE sepharose (Figure 1B). This was followed by ultracentrifugation through a sucrose gradient to separate the fragments by length to further remove contaminating proteins and free DNA. By adjusting the amount of MNase and the conditions of ultracentrifugation it was possible to fine-tune the proportions of nucleosomal species, ranging in length from mono- to tetra-nucleosomes (Figure 1C). Under such limiting conditions, little or no over-digestion of nucleosomes occurred. Indeed, these mono-nucleosomes were similar in nature to nucleosomes previously defined as ‘under-digested’ by (Weiner et al. 2010). The final material used consists entirely of mono-nucleosomes and is of high purity, as judged by SDS-PAGE analysis (Figures 1C and D). *In vitro* ChIP of selected histone marks across a previously characterized gene (Kim and Buratowski 2009) showed the expected profile (Figure 1E), indicating that the epigenetic marks of the original material are indeed maintained. Paired-end sequencing of purified mono-nucleosomes followed by mapping of the reads to the yeast genome showed the typical nucleosomal pattern (Figure 1F; see also below and Figure S2E). The majority of the genome was recovered in the purification: 94% was thus covered by at least 5 reads. As might be expected, areas with less, or no, reads were mainly found towards the gene-poor ends of the chromosomes (Figure S1). This agrees with the previous finding that yeast chromatin is mostly open and active (Rattner et al. 1982) and shows that native nucleosomes are generally stable enough to withstand stringent purification.

**Figure 1.**
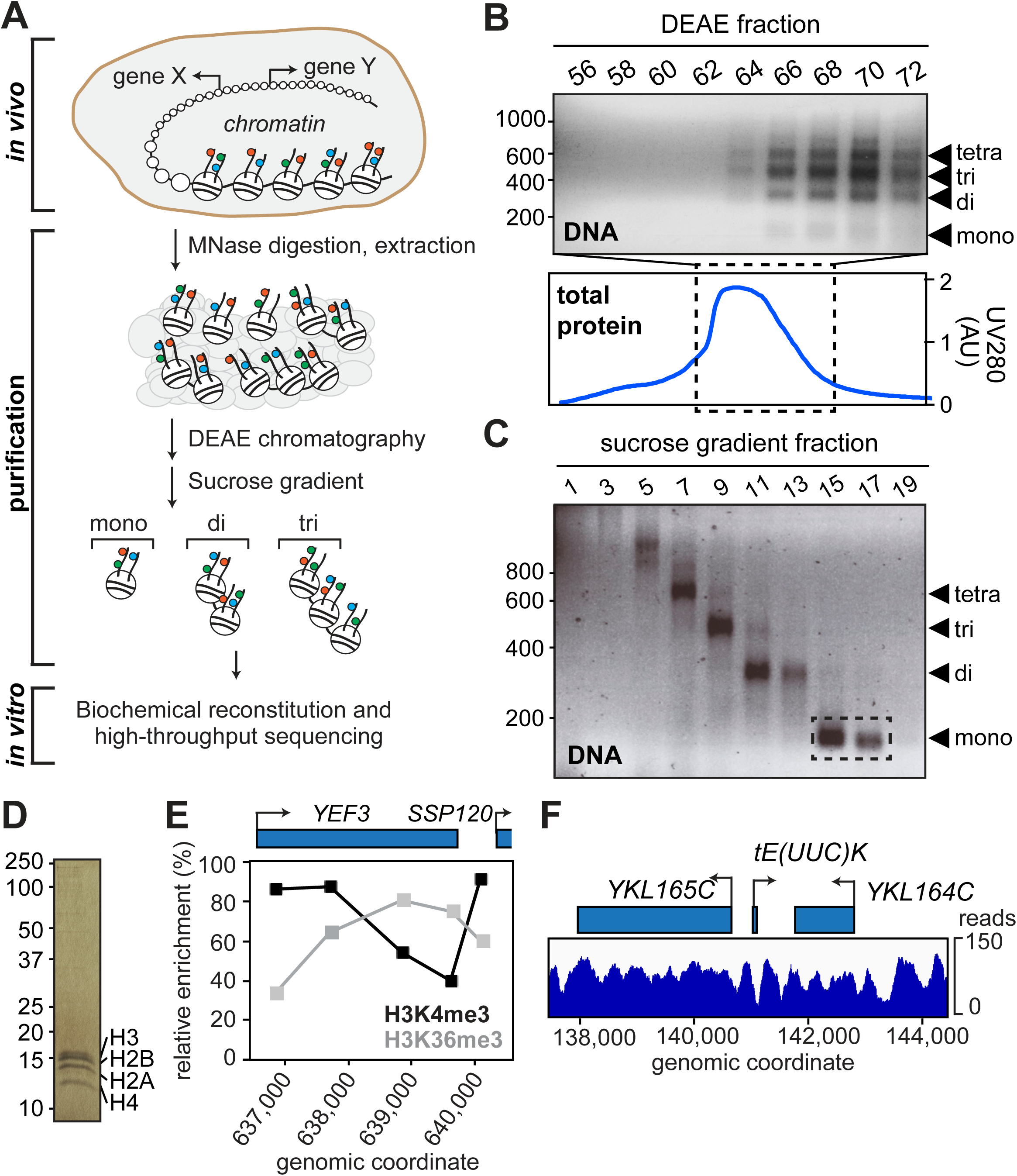
Purification and characterization of genomic chromatin. **A.** Schematic of experimental approach. Colored circles, histone marks. **B.** DNA from DEAE fractions analyzed by agarose gel-electrophoresis (top panel); marker on left shows length in bp. Bottom panel, chromatogram of total eluted protein. Fractions 66-72 were loaded on the sucrose gradient. **C.** DNA from sucrose gradient analyzed by agarose gel-electrophoresis. Fractions used for experiments are indicated by stippled box. **D.** Silver-staining of purified mono-nucleosomes from C. **E.** Histone mark patterns of purified mono-nucleosomes as determined by native ChIP-qPCR. Histone marks were normalized to histone H3, as in the reference datasets of Pokholok *et al.* (2005) and Kim *et al*. (2009). **F.** Representative map of nucleosomes on chromosome XI after paired-end sequencing and alignment to the yeast genome.

A number of assays for measuring chromatin remodeling *in vitro* have been developed. We chose a simple disassembly assay, which involves incubating the nucleosome library with ATP and the histone chaperone Nap1, with or without RSC (Lorch et al. 2006). In this assay, RSC binds to nucleosomes and transfers the histones to Nap1, thereby releasing ‘naked’ DNA (Figure 2A). Under certain conditions, reaction intermediates can be observed (tetramers or hexasomes (Lorch et al. 2006; Kuryan et al. 2012), but for simplicity we chose to compare the input nucleosomes with the final naked DNA product. While it represents a further simplification, the general term ‘remodeling’ is hereafter used to describe the successful RSC-dependent release of such products. To separate the ejected DNA product from the non-remodeled nucleosomes, the reactions were subjected to native agarose gel electrophoresis (Figure 2B), and DNA of the four bands isolated by gel-extraction. The upper bands, harboring nucleosomes, were named NUC (no RSC) and NUCR (with RSC), whereas the lower, ‘naked’ DNA bands were named DNA (no RSC) and DNAR (with RSC). A set of control reactions for the RSC-dependent reaction confirmed that the assay was indeed dependent on Nap1, ATP and RSC (Figure S2A). Over limiting time, RSC remodels only a subset of the input nucleosomes (Figure S2B, left). Although we here confined our analysis to mono-nucleosomes, the reaction also worked on di-nucleosomes (Figure S2B, right). The extracted DNA was sequenced after paired-end adapter ligation, enabling us to map each nucleosome to the reference yeast genome. Fragments from all four bands had the length expected for nucleosomal DNA containing a short linker (Figure S2C). Indeed, no dramatic differences in the mean length of nucleosome fragments were observed between the four different categories (NUC, NUCR, DNA, DNAR), or across the different stability groups defined below (Figure S2D). Even after experimentation and gel-extraction of the underlying DNA (Figure 2A and B), the majority of the genome was still detected by deep sequencing of the DNA libraries: over 83% (80%) of the genome was thus covered by at least 5 reads in DNA + NUC (DNAR + NUCR) in at least one of the experimental replicates, and both replicates again showed the typical nucleosomal pattern (Figure S2E). These patterns are highly similar to those previously reported by Kaplan *et al* (2009), also using MNase digestion. The pile-ups of reads across bands after incubation showed that these indeed came from the same individual nucleosomes (compare pile-ups of reads in DNA and NUC in the example shown in Figure 2C).

**Figure 2.**
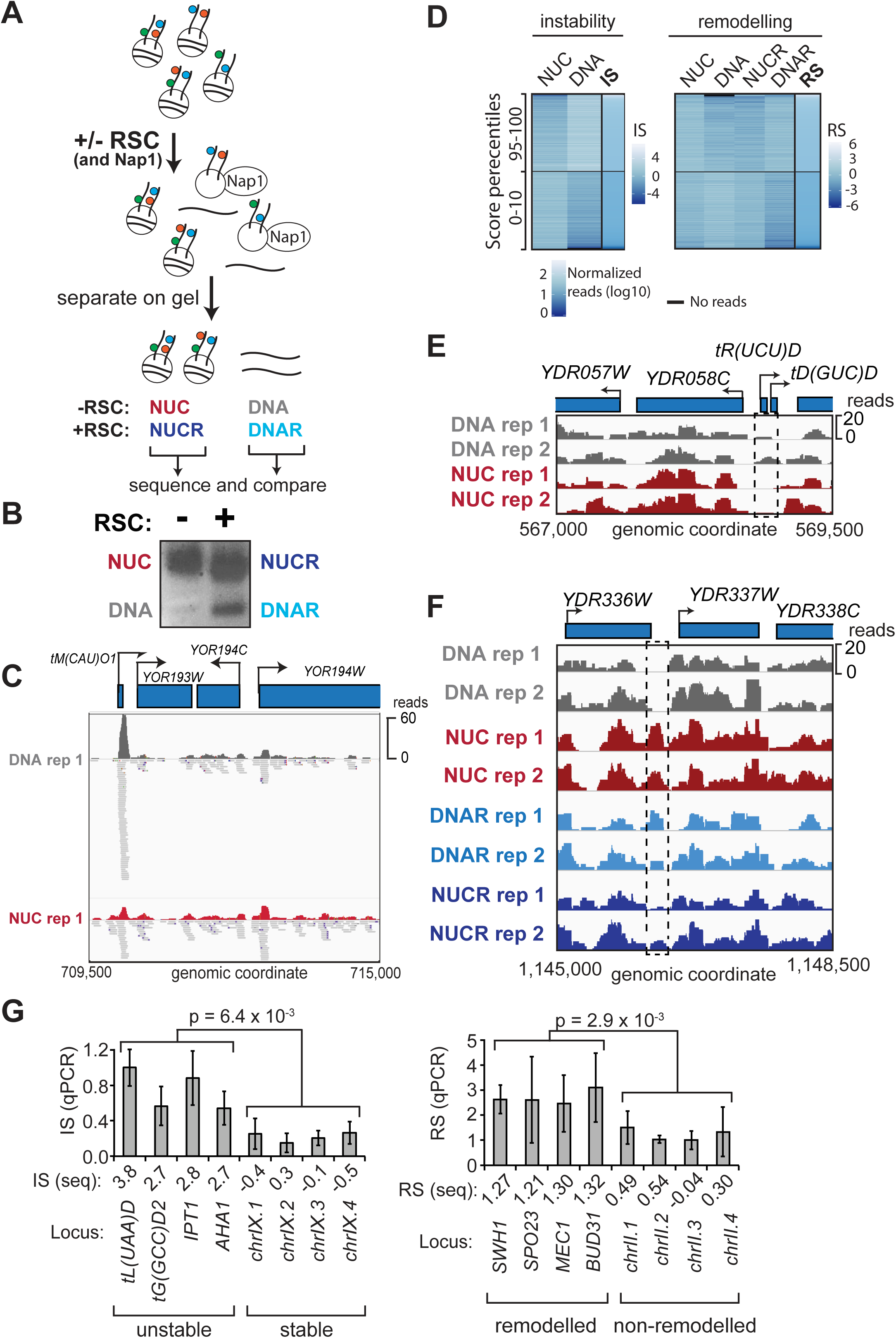
Chromatin remodeling and instability studies performed on native chromatin. **A.** Schematic of the nucleosome disassembly assay. **B.** Nucleosomes incubated +/-RSC were separated by native agarose gel electrophoresis. The four indicated bands were excised and the DNA extracted and sequenced. **C.** Representative example of highly unstable nucleosome at tM(CAU)O1 (chromosome XV), enriched in the ‘DNA’ band after incubation, but also detected in ‘NUC’. **D.** Heatmaps of read densities for nucleosomes with different IS and RS, respectively, comparing distinct percentiles. **Left** boxes, distribution of read densities in bands NUC and DNA for corresponding IS distributions. **Right** box, equivalent distribution for bands used to calculate RS. 10,000 randomly sampled windows in the 10th and 95th percentiles are shown. **E.** Region encompassing a representative, highly unstable nucleosome (dashed box; chromosome IV (IS =1.0). **F.** As E, but for a strongly remodeled nucleosome (chromosome IV (RS =0.78)). **G.** qPCR validation, using primers for the indicated regions. **Left,** plot of instability index for qPCR (IS-qPCR). Four unstable nucleosomes followed by four stable nucleosomes. The p-value calculated using the Wilcoxon t-test; error bars show standard deviations from four biological replicates. **Right,** as left, but for remodeling. Three biological replicates were performed.

Nucleosome position calling from MNase-derived sequencing data is very dependent on the specific software, so we chose a window-based approach as a robust method for quantifying nucleosome occupancy. For this purpose, we divided the yeast genome into 167 bp sliding windows with a step size of 25 bp. As previously reported for MNase-Seq for nucleosomal DNA (Chung et al. 2010), we observed some enrichment for GC-rich reads and therefore normalized the read counts for the GC content for each. After normalization and quality filtering, 450,257 windows were recovered for comparison.

In order to quantify the intrinsic stability of each nucleosome, we calculated the normalized log-ratio of reads between the DNA and NUC samples per window using both replicates. The resulting Instability Score (IS = log_2_(DNA/NUC)) measures the relative stability of nucleosomes across the genome, with high scores indicating more unstable nucleosomes. Similarly, we quantified the degree of RSC-dependent nucleosome disassembly by calculating the log-ratio of reads from the DNAR and NUCR samples, before subtracting the Instability Score to account for intrinsic instability. The resulting Remodeling Score (RS = log_2_(DNAR/NUCR) – IS) measures the effect of RSC-dependent remodeling, with higher scores reflecting a more strongly remodeled nucleosome. We divided the sets in 8 percentiles (0-10%, 10-25%, 25-50%, 50-75%, 75-90%, 90-95%, 95-99%, and 99-100%). Finally, overlapping or adjacent windows in the same IS/RS-percentile were merged to obtain continuous unstable or remodelled regions. For simplicity, we will hereafter often refer simply to ‘nucleosomes’ although these are strictly speaking merged, overlapping windows in the same IS/RS-percentile. We hereby obtained sets of 343,860 windows (corresponding to 82,663 nucleosomes) to study instability, and 229,479 windows (66,628 nucleosomes) to study RSC remodelling, respectively.

Figure 2D shows a heatmap of read counts in the four bands and how they relate to the scores. The higher the IS, the more naked DNA there is compared to nucleosomal DNA (DNA/NUC). For RSC-dependent remodeling, not only must nucleosomes display higher read counts in the DNAR sample compared with NUCR, but they must also be intrinsically stable in order to achieve a high RS. In general, the IS and RSs are thus anti-correlated. It is worth noting that while the RSs can, of course, be compared amongst each other, their actual value do not have an intuitive meaning (as opposed to the IS), and even ‘high’ RS are frequently negative due to the normalization with the IS. For much of the remainder of the analysis we often focus on the 99^th^ percentile, which comprises the ∼3,400 most intrinsically unstable windows and the ∼2,300 most remodeled windows, corresponding to ∼1100 and ∼900 nucleosomes, respectively. Figures 2C and 2E shows a representative intrinsically unstable nucleosome (stippled boxes), and Figure 2F shows a representative strongly remodeled nucleosome. For validation, we performed qPCR analyses on eight representative stable/unstable and remodeled/non-remodeled nucleosome positions (Figures 2G).

### Reduced intrinsic stability of tRNA and RNAPII promoter nucleosomes

Interestingly, the nucleosomes represented by the different instability percentiles were not randomly distributed across the genome. Instead, the most unstable nucleosomes were mainly found overlapping with protein-coding genes, tRNA genes, and promoters (Figure 3A and S3A). Given the high instability at tRNA genes, we further analyzed their IS. At such genes, the IS was low in the area from −600 to −200 (suggesting stable nucleosomes in this area), and then increased drastically to reach maximum instability directly over the Pol III-transcribed region (suggesting a highly unstable nucleosome) (Figure 3B, left). However, these distinct nucleosome types do not co-exist on the same gene; instead, there is a subset of tRNA genes (in the 10^th^ percentile) that had a very stable nucleosome(s) upstream from the transcribed region, but these genes are separate from the group of tRNA genes (in the 99^th^ percentile) that had a highly unstable nucleosome directly on the transcribed region (Figure 3B, right). Unlike protein-coding genes, which are occupied by multiple nucleosomes, the short tRNA genes are typically covered by only a single nucleosome, which thus appears to be intrinsically highly unstable. 89 of the 275 yeast tRNA genes contained a highly unstable nucleosome (99^th^ percentile) and tRNA genes were generally highly enriched in nucleosomes in the 95^th^ percentile (Figure S3B); nucleosomes overlapping with tRNA gene bodies thus generally had substantially higher IS compared with all others (p-value <0.01, 2-sided Wilcoxon Rank Sum test).

**Figure 3.**
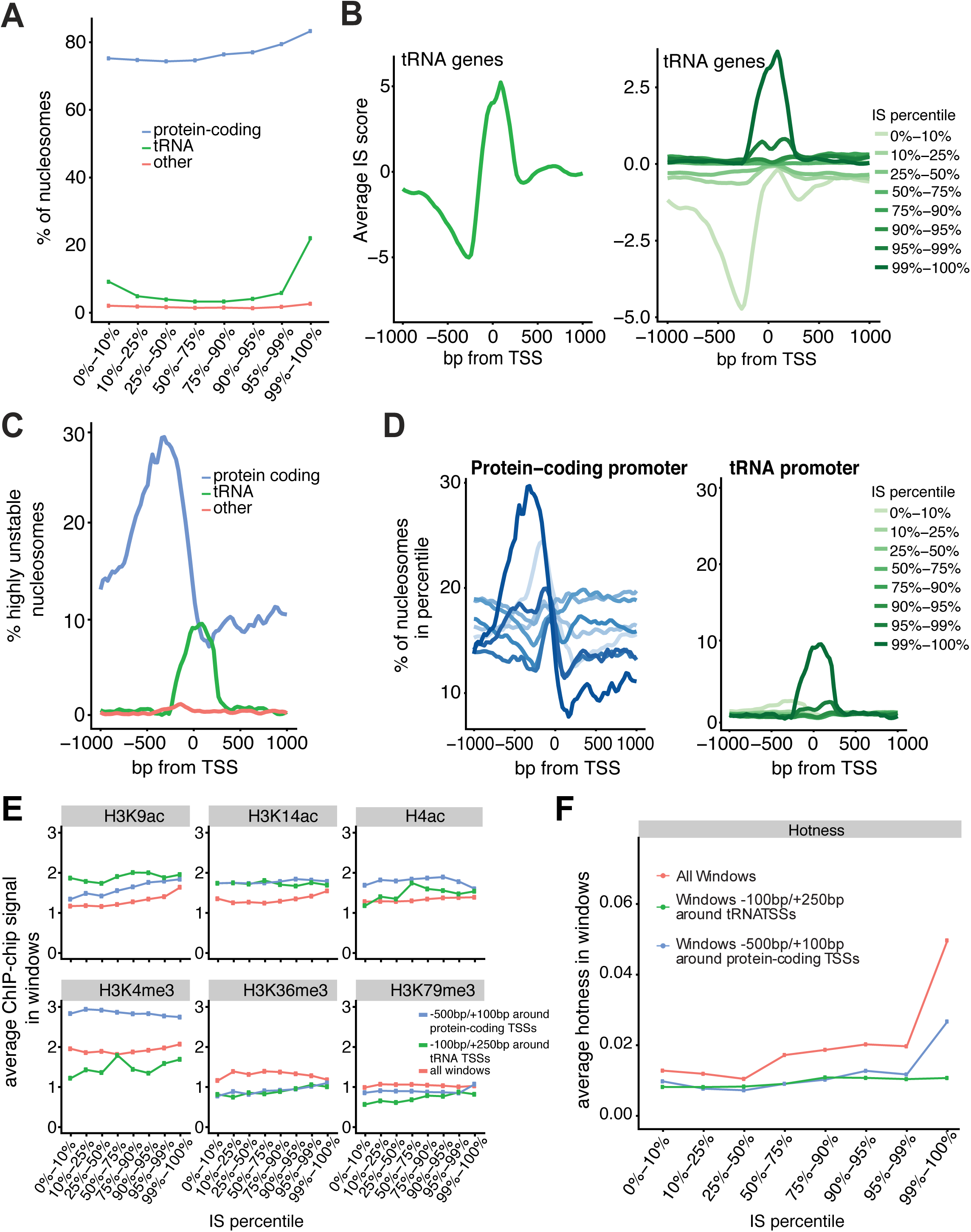
Characterization of intrinsically unstable nucleosomes. **A.** IS percentiles plotted against nucleosomes that overlap with genes. **B. Left**, average ISs around the TSS of tRNA genes. **Right,** as left, but averages observed in different IS percentiles. **C.** Position around the TSS of nucleosomes in the 99^th^ IS percentile. **D. Left,** as C, but nucleosomes in different IS percentiles around the TSS of protein-coding genes. **Right**, as left, but for tRNA. **E.** Histone marks in percentiles, for nucleosome windows in different genomic contexts. **F.** Hotness of nucleosome windows in percentiles, around genes and across the genome.

We also observed an enrichment of protein-coding gene promoters among nucleosomes in the higher percentiles (Figure S3C). Plotting the position of the nucleosomes in the 99^th^ percentile, a broad area of intrinsic instability was uncovered, peaking in the promoter upstream of protein-coding genes (Figure 3C). In contrast to the peak representing tRNA genes, which was only observed with very high ISs (Figure 3D, right), a peak in the promoter of protein-coding genes was seen in the highest percentiles, but also in the lowest percentile (Figure 3D, left), suggesting that both highly unstable and highly stable nucleosomes were detected in this area, depending on the gene. qPCR analysis showed that the release of DNA from unstable nucleosomes was indeed independent of ATP and Nap1 (Figure S3D), and instability could therefore be classified as intrinsic to the structure of the nucleosome.

We sought to better understand the nature of the intrinsically unstable nucleosomes and even considered the possibility that these might not actually be nucleosomes to start with, but – for example - fragments that were cut out by MNase as naked DNA and ‘contaminated’ the input material. Several observations indicated that this was not the case. First, the input nucleosomes were not merely isolated by sucrose gradient purification, but also by DEAE chromatography, with elution by a salt gradient. Mono-nucleosomes elute at ∼400 mM NaCl, whereas free DNA elutes much later (Luger et al. 1999). Free DNA should thus not be present in the input nucleosome sample. Second, we compared the nucleosomes in the different IS percentiles with published nucleosome datasets measured by MNase-Seq (Kaplan et al. 2009), or chemical cleavage (Brogaard et al. 2012). Although a slight decrease in number of nucleosomes detected by MNase digestion or chemical cleavage was observed in the windows representing the very highest ISs, nucleosomes clearly exist in these regions (Figure S3E, upper and lower, respectively). Actually, in the vast majority of positions with highly unstable nucleosomes (92%, N=1,112) nucleosomes were also detected by Widom and co-workers by chemical cleavage (Brogaard et al. 2012). Third, naked DNA fragments cut out by MNase might be expected to differ in size from true nucleosomes, yet the DNA reads that mapped to tRNA gene bodies were highly similar in length to those of the total pool of DNA reads (Figure S3F), with no statistically significant difference detected (2-sided Wilcoxon Rank Sum test). Similarly, there were no differences in the lengths of DNA reads across the different stability groups, nor between DNA and NUC reads (Figure S2D). Finally, visual inspection of the mapped read files shows that there is nothing unusual about the reads representing these nucleosomes: most importantly, they *were* also detected among the <u>gel-purified</u> mono-nucleosomes but were indeed enriched in the free DNA sample after incubation (See example in Figure 2C). We conclude that many nucleosomes on tRNA genes and on promoters of protein-coding genes are intrinsically highly unstable.

We next examined whether any known genomic features might explain the results. GC content influences nucleosome occupancy *in vivo* and *in vitro* (Tillo and Hughes 2009), and poly(dA:dT) tracts decrease nucleosome assembly *in vitro* (Anderson and Widom 2001). However, GC-content (Figure S4A) was only slightly decreased, and poly(dA:dT) tracts were slightly <u>in</u>creased in the windows with the highest IS percentiles (Figure S4B). Intriguingly, while the IS reached its maximum at the dyad of the highly unstable tRNA nucleosomes, the dyads were actually AT-depleted (Figure S4C). The lack of a clear correlation with these sequence characteristics (low GC content and AT-tracts) also distinguishes these intrinsically unstable nucleosomes from the unstable/’fragile’ nucleosomes studied by, for example, the Shore laboratory (Kubik et al. 2015).

We also examined whether unstable nucleosome positions coincided with specific histone modifications, using published ChIP-chip datasets for three methylation marks (H3K4me3, H3K36me3 and H3K79me3) and three acetylation marks (H3K9ac, H3K14ac and H4ac) (Pokholok et al. 2005). H3K9 and H3K14 acetylation is generally enriched around the TSSs of both tRNA and protein-coding genes compared with all genome-wide windows (Figure 3E). While the enrichment of H3K14 acetylation was similar across all IS percentiles in both protein-coding and tRNA genes, there was a tendency for H3K9 acetylation to increase slightly with increasing IS, particularly around the TSSs of protein-coding genes. Similar, but less obvious trendlines were also observed for other marks, but only increasing very modestly from relatively low levels (Figure 3E; see H3K4me3, H3K36me3, and H3K76me3 for tRNA genes, for example). Interestingly, histone H2A.Z was depleted from highly unstable nucleosomes (Figure S4D).

We did not find a correlation between the ISs of promoter regions or TSSs and the transcription levels of the associated genes, as measured by RNA-seq (Nagalakshmi et al. 2008) (data not shown). However, promoters of genes have previously been reported to preferentially be occupied by “hot” nucleosomes; nucleosomes that for unknown reasons are exchanged rapidly outside of S-phase *in vivo* (Dion et al. 2007). We found that while protein-coding promoters indeed had a high level of ‘hotness’, there was a clear tendency for the highly unstable windows to be hotter in general (Figure 3F, protein-coding promoters and all windows).

We conclude that many nucleosomes on tRNA genes and on promoters of protein-coding genes are intrinsically unstable, but that this is not generally correlated with DNA sequence features previously found to affect nucleosome behavior. Instead, intrinsic instability correlates weakly with a modest increase in a variety of histone marks (Pokholok et al., 2005), including H3K9Ac, H4Ac and H3K4me3. The finding that high ISs correlated with high levels of histone exchange *in vivo* (i.e. hotness) may help explain the underlying mechanism of hotness: many such highly exchanged nucleosomes are likely intrinsically unstable.

### Preferential disassembly of NFR- and TSS nucleosomes by RSC

Next, we investigated the characteristics of the nucleosomes that were remodeled by the RSC chromatin remodeler. As observed for intrinsically unstable nucleosomes, the nucleosomes represented in the different remodeling percentiles were not randomly distributed across the genome. Indeed, the most strongly remodeled nucleosomes were relatively enriched in gene promoters, and in so-called nucleosome-free regions (NFRs, defined as 250bp-50bp upstream of the TSS) (Figures S5A and 4A). In contrast to the Instability Scores, only minor differences between protein-coding and tRNA genes were uncovered for Remodeling Scores, so for simplicity, the analysis below is focused on mRNA and tRNA genes together (‘genes’). Analysis of the RS within 1 kb of the TSS thus showed a marked increase of the score just upstream of, and on, the TSS (Figure 4B). Importantly, these highly remodeled nucleosomes were not the same as the highly intrinsically unstable nucleosomes: indeed, only 6 windows genome-wide were classified as both highly unstable *and* strongly remodeled. In fact, as mentioned above, due to the normalization of the RS by the IS, these scores are generally anti-correlated.

**Figure 4.**
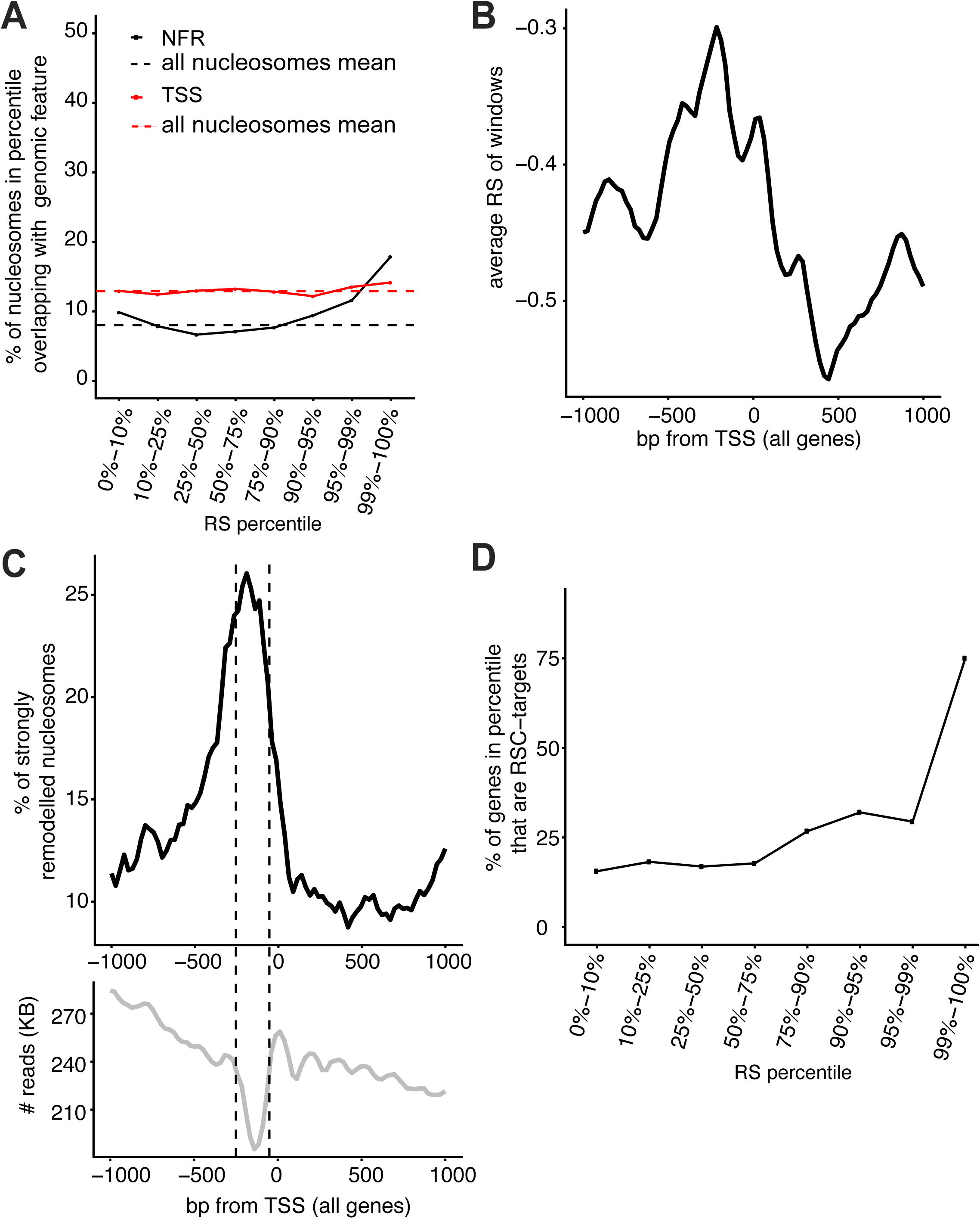
Characterization of nucleosomes remodeled by RSC. **A.** Distribution of nucleosomes in different RS percentiles that overlap with the TSS or NFR (50-250bp upstream of TSS). **B.** Average RS around the TSS of genes. **C.** Upper, Metaprofile around the TSS of genes of strongly remodeled nucleosomes (99^th^ percentile), relative to the general nucleosome density in the same region (lower graph). **D.** RSC target genes in the different RS percentiles, relative to total number of genes in same.

When we aligned the most strongly remodeled nucleosomes (99^th^ percentile; ∼900 nucleosomes) around the TSS’s of genes, we found a broad peak upstream of the TSS, covering both the n-1 and n+1 nucleosome, and peaking in the NFR (Figure 4C). Of the most strongly remodeled windows, 19% were in the NFRs (indicated by dashed lines in Figure 4C, lower), compared with 9% of all windows. As an important control, we investigated the lengths of the nucleosome fragments in the DNAR vs NUCR samples (Figure S2D). The lack of significant size differences between the different percentiles indicated that the highly remodeled nucleosomes do not carry additional DNA, such as adjacent promoter DNA that might have served to recruit RSC, and that the recruitment signal(s) must thus be contained within the nucleosome itself.

We conclude that RSC prefers to remodel mono-nucleosomes derived from promoter and TSS regions, and particularly from the so-called nucleosome-free regions.

### RSC preferentially disassembles nucleosomes originating near genes it regulates

What characterizes nucleosomes that are most strongly remodeled by RSC? GC-content did not appear to be a major defining variable although there was a modest decrease in the higher RS percentiles (Figure S5B). Poly(dAdT) tracts were generally modestly enriched in remodeled nucleosomes (Figure S5C), in apparent agreement with the previous finding that RSC shows a preference for reconstituted nucleosomes bearing poly(dAdT) tracts (Lorch et al. 2014).

RSC harbours eight bromo-domains, at least one of which has been shown to bind acetylated histone tails (Kasten et al. 2004). Somewhat to our surprise, we failed to find one or more histone marks that were markedly enriched at higher RSs (Figure S5D). As observed with the IS, we also failed to find a general correlation between RS and the transcription levels of the corresponding genes across the genome, as measured by RNA-seq (Nagalakshmi et al. 2008). Importantly, however, we also investigated possible connections between the RS at the TSS and transcription of genes that were previously found to be dependent on RSC for their expression (Parnell et al. 2015). Gratifyingly, when the 615 genes tested by Parnell *et al.* were sorted into the different RS percentiles, the 124 genes affected by RSC were enriched at high RS (Figure 4D), with more than a third of the genes in the 90^th^ percentile being RSC-dependent genes, and RSC target genes generally having a higher RS than non-targets (p-value <0.01, one-sided Wilcoxon rank sum test). This suggests that the characteristics of chromatin at these genes are preserved to some degree in isolated mono-nucleosomes, and, vice versa, that recognition of individual nucleosomes by RSC *in vivo* may indeed have significant consequences for transcription of the adjoining gene.

### RSC preferentially disassembles H2A.Z-containing nucleosomes

Arguably, DNA sequence and histone marks provided somewhat limited information about the mechanism underlying the nucleosome preference of RSC. However, another candidate feature is the histone variant H2A.Z, which is enriched around the TSS of genes (Albert et al. 2007), in a profile which is very similar to that of the highly remodeled nucleosomes (compare Figure S6A with Figure 4B). We therefore looked specifically at nucleosomes containing H2A.Z. For this purpose, we categorized nucleosomes as H2A.Z+ if they were in the top 30^th^ percentile of H2A.Z levels (Albert et al. 2007). Interestingly, many of the strongly remodeled nucleosomes making up the peak just upstream of the TSS indeed carried H2A.Z (Figure 5A). Moreover, the proportion of promoter regions carrying H2A.Z increased with the RS of those regions (Figure 5B). Promoters that carry H2A.Z have been suggested to preferentially be occupied by nucleosomes that are rapidly exchanged *in vivo* (“hot” nucleosomes; (Dion et al. 2007)). Interestingly, however, the RS was not correlated with ‘hot’ nucleosomes near the 5’-end of genes (Figure S6B), in contrast to the IS (see above and Figure 3F). This indicates that the efficient RSC-mediated remodeling of H2A.Z-containing nucleosomes observed *in vitro* is not due to such nucleosomes being particularly unstable or exchanged rapidly.

**Figure 5.**
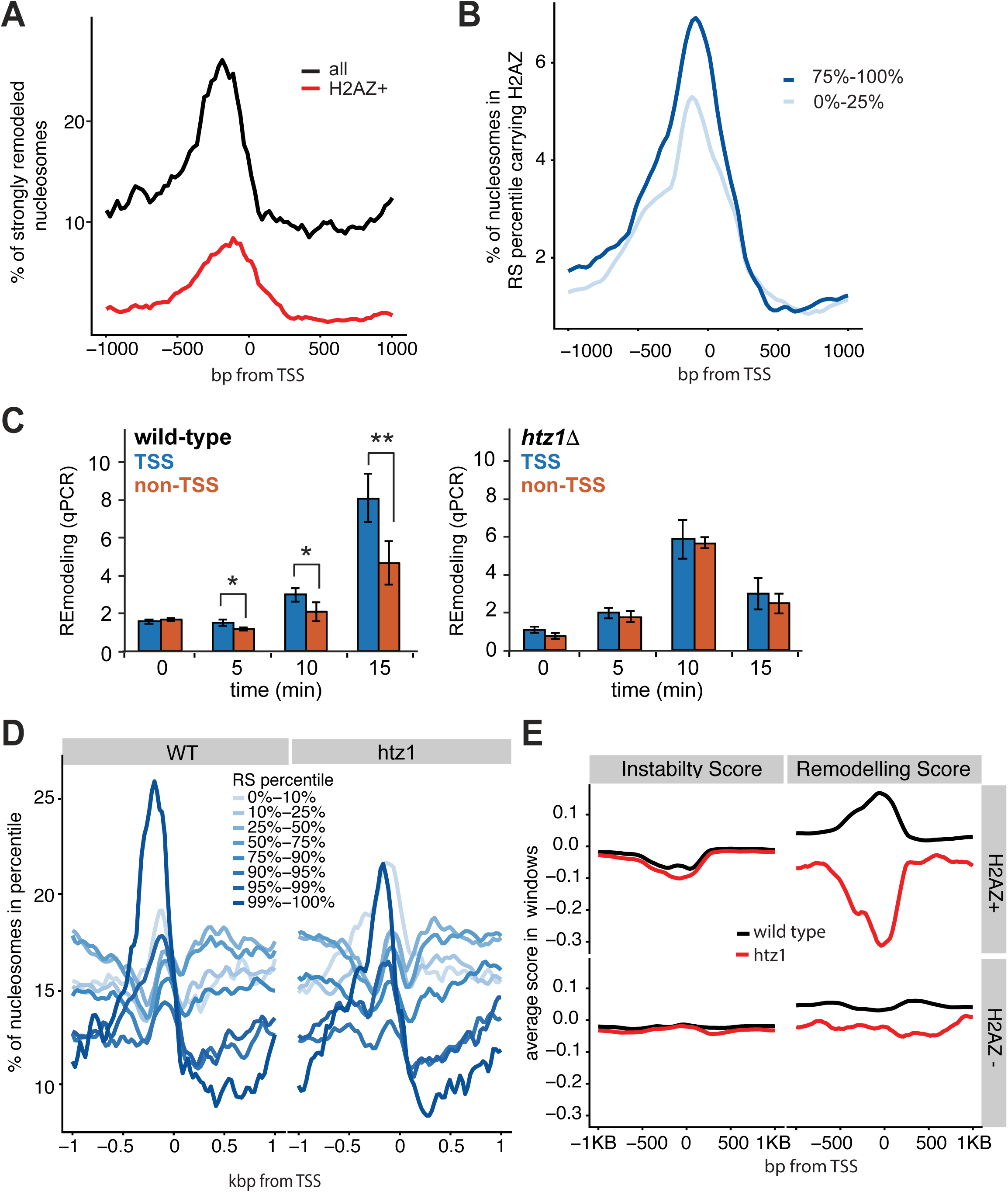
RSC prefers H2A.Z-containing nucleosomes. **A.** H2A.Z-containing nucleosomes in the 99^th^ RS percentile and their position around the TSS (red), compared with pattern for all nucleosomes in same (black). **B.** H2A.Z as in A, but in different RS percentiles. **C.** Time-course of RSC remodeling preference with nucleosomes from wild-type (left) and *htz1*Δ, (right) by qPCR. Blue bars show average of four strongly remodeled, H2A.Z+ TSS nucleosomes, and orange bars show average of four control nucleosomes (see Figures S6D and E). Asterisks show statistical significance (* = p < 0.05; ** = p < 0.01; Student’s t-test). **D.** Position around the TSS of nucleosomes in the different RS percentiles, for wild-type (WT) and *htz1*Δ nucleosomes, respectively. **E.** Comparison of the IS and RS for H2A.Z+ and H2A.Z-around the TSS of genes. For ease of comparison, plots were scaled individually to range between −1 and +1 genome-wide.

In order to experimentally address a putative causative role for H2A.Z in the preference of RSC for TSS nucleosomes, we performed remodeling assays using nucleosomes prepared from *htz1*Δ cells (Figure S6C) and compared the resulting preference with that observed for wild type nucleosomes. Initially, we selected four individual nucleosomes that we had found to be strongly remodeled and which carry H2A.Z (Albert et al. 2007). These were compared with four control nucleosomes (Figure S6D). Using qPCR to detect remodeling in these regions, we found that RSC indeed showed a preference for H2A.Z-containing nucleosomes at the TSS when presented with nucleosomes from wild type cells, but not with nucleosomes from *htz1*Δ cells, even though htz1Δ nucleosomes were, of course, still remodeled due to the general ‘background’ activity of RSC (Figure 5C and S6E).

We also prepared DNA libraries from experiments with *htz1*Δ nucleosomes for deep sequencing and analyzed the resulting data in the same manner as before, computing Instability- and Remodeling Scores genome-wide. Parameters such as genome coverage, nucleosome positioning, nucleosome fragment lengths, and the distribution of raw DNA and NUC reads to windows in the different IS percentiles were similar to those observed in wild type, indicating that the data-sets were suitable for comparison (Figure S2C, S7A and B; and data not shown). The nucleosome occupancy metaprofiles in the different RS percentiles were similar between WT and *htz1*Δ nucleosomes around the TSS of genes, with some notable exceptions: the large peak of the 99^th^ percentile in WT was much reduced with *htz1*Δ nucleosomes and the 95^th^ percentile peak was largely absent (Figure 5D). This was accompanied by the 10th percentile for *htz1*Δ showing a peak of the same height as that of the 99th percentile. Indeed, only 10% of the strongly remodeled windows (99^th^ percentile) in WT were also detected among in the 90^th^ percentile RS with *htz1*Δ nucleosomes, and a sub-group from the 99^th^ percentile in WT of almost the same size (9%) moved to the bottom 10th percentile of the *htz1*Δ RS. This indicates an underlying re-ordering of the preferred nucleosome remodeling positions in *htz1*Δ cells.

To further analyze these data, we again divided all windows into H2A.Z+ (as above; top 30^th^ percentile H2A.Z density (Albert et al. 2007)) and H2A.Z- (bottom 30^th^ percentile)) and compared their IS and RS around the TSS of genes (Figure 5E). Remarkably, while the meta-profiles for the IS and RS for H2A.Z-windows were similar (lower panels), the meta-profiles of the RS of the H2A.Z+ windows displayed a stark difference between WT and *htz1*Δ (upper panel on the right): the WT RS thus increased on the TSS, while the opposite was observed with the *htz1*Δ RS, which reached a minimum in the same area. Importantly, the ISs for the same regions were similar so cannot account for this difference (Figure 5E, upper panel on the left). Given the strong effect of H2A.Z, we also attempted to uncover any (previously hidden) characteristics of nucleosomes that are highly remodeled (in the 90^th^ percentile) in *both* WT and *htz1*Δ, but failed to find any enrichment in sequence features (GC-content or poly(dA:dT)-tracts), and only a modest, and counter-intuitive, depletion in H4 acetylation levels (data not shown). We conclude that RSC prefers to remodel mono-nucleosomes originating from around the TSS in genes and that this preference is to a significant degree mediated via H2A.Z.

To finally investigate nucleosome remodeling by RSC using an independent, complementary system, we employed nucleosome arrays that were assembled on circular DNA plasmids using recombinant histone proteins in the presence of DNA topoisomerase I (Clapier et al. 2016)(Figure 6A). Arrays assembled with either H2A or H2A.Z-containing nucleosomes showed similar topoisomer distribution (Figure 6B, left). Nucleosome ejection by RSC alone from such plasmids causes a change in topoisomer distribution, with progressively less assembled states (and successively fewer nucleosomes) distributed in a clockwise manner along an arc, and any unassembled plasmids present at the lower right terminus (Clapier et al. 2016)(Figure 6A). Gratifyingly, RSC was much more efficient at nucleosome ejection with the H2A.Z arrays than with arrays assembled with canonical histone H2A (Figure 6B, right). These results strongly support the finding that RSC preferentially ejects nucleosomes containing H2A.Z.

**Figure 6.**
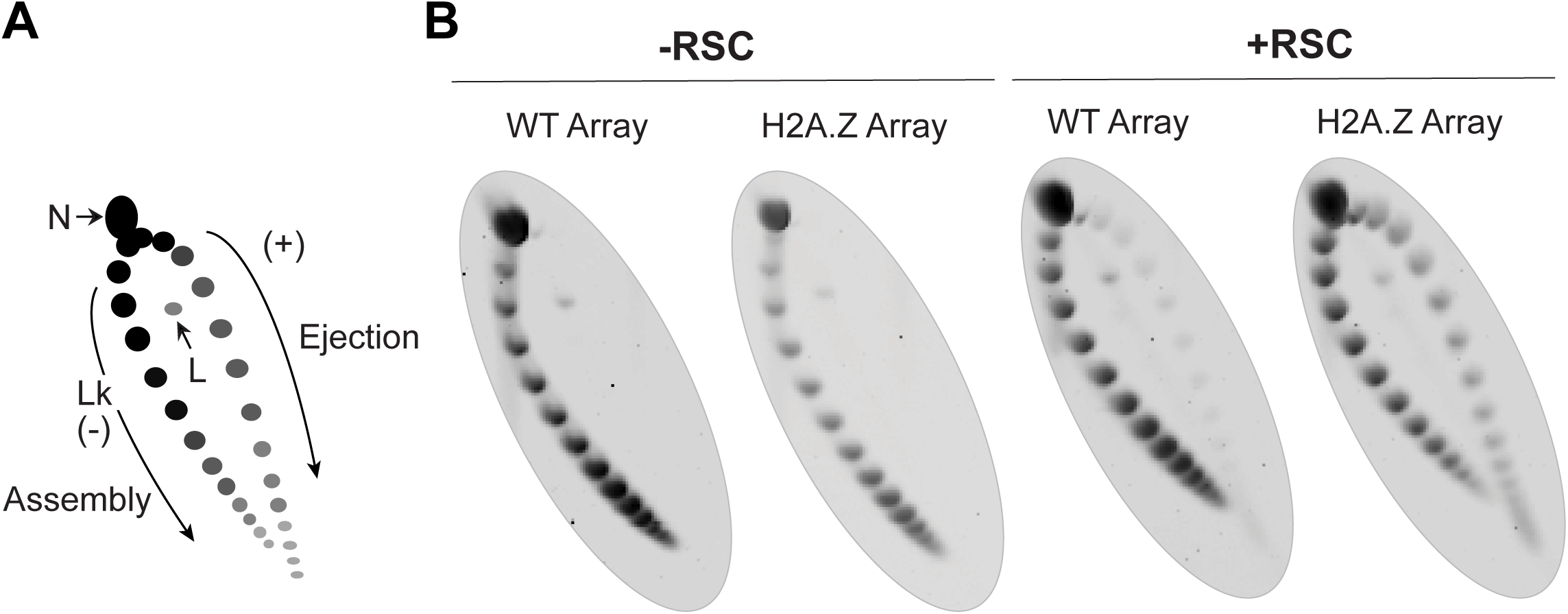
Preferential ejection of H2A.Z-containing nucleosomes from arrays by RSC. **A.** Schematic of the principle of the nucleosome array ejection assay, with supercoiled plasmid (topoisomer) distribution revealed by 2D gel. Lk: Linking number, N: Nicked, L: Linear. **B.** Nucleosome arrays assembled with canonical octamers (WT Array) or with H2A.Z-containing octamers (H2A.Z Array), incubated +/-RSC.

## DISCUSSION

The challenge of understanding the processes that take place in natural, eukaryotic chromatin, particularly at the molecular or biochemical level, is substantial. Reconstituting such chromatin with purified components *in vitro* is for obvious reasons extremely challenging, and the study of biochemical mechanisms *in vivo* is very difficult as well. Here, we present a new molecular tool, which we believe can help fill the gap between these approaches: a ‘library’ of mono-nucleosomes, which can be used to study different biochemical characteristics of natural chromatin, as well as the substrate preference of chromatin binding factors, ATP-dependent remodelers, and chromatin-modifying enzymes. The validity of the approach is suggested by the results themselves: the finding that both intrinsically unstable nucleosomes and RSC-remodeled nucleosomes have specific and highly sensible characteristics (i.e. genome position, underlying DNA sequence, and H2A.Z content, for example) indicates that the methodology works. One might have feared, for example, that random nucleosome regions had resulted from these experiments. Instead, the nucleosome-selecting factor tested – the chromatin remodeler RSC – does indeed make meaningful choices from this library, which help further the understanding of the biology of both nucleosomes, chromatin, and RSC.

What specifies a particular genomic location - whether it is a promoter, a TSS, a telomere, or a centromere? In many cases, there are conserved DNA sequence motifs, as is the case with telomeres and recognition-sites for DNA-binding transcription factors, for example. But what when there are no conserved sequence elements, as is the case for many promoters and TSSs? One obvious possibility is that it is exclusively through the combined presence of nearby protein-binding sites that a genomic locus comes to acquire its functional identity. According to that model, the spatial context in which a region exists is crucial for that area to ‘know’ its function. Some evidence for this idea has grown with the advent of techniques for mapping long-range interactions on chromatin (de Wit and de Laat 2012; Denker and de Laat 2016). However, even though distant interactions affect local function, they are unlikely sufficient to establish local identity. For one, exclusive dependence on distal regions would severely restrict co-evolution of functionally linked areas. It would also fail to explain how regions of short length can often be cloned into another genomic locus, while apparently remaining fully functional. It thus seems likely that loci also carry information about their identity on a local scale. The results of this study indicate that, actually, a surprisingly large amount of information about regional genomic function may be inherent, contained in single nucleosomes, and independent of the surrounding context. This applies to at least three classes of nucleosomes: unstable nucleosomes on promoters of protein-coding genes and on tRNA genes, and nucleosomes near the TSS of genes, the latter of which were found to be preferential targets of RSC. Crucially, identity is preserved in the individual nucleosome, free from the necessity for any higher-order chromatin structure: when isolated and incubated *in vitro*, these mono-nucleosomes contain sufficient information to continue to at least partially function as if they were still embedded in their natural genomic context.

Previous work has shown that RSC plays a crucial role in establishing NFR’s *in vivo* (Hartley and Madhani 2009) and *in vitro* (Wippo et al. 2011). Using mono-nucleosomes purified from cells, we now find that RSC preferentially remodels the individual mono-nucleosomes on the TSSs and promoter regions, and particularly those within the NFR. This complements, generalizes, and extends the discovery that RSC intrinsically prefers the promoter nucleosomes on a purified *PHO5* chromatin circle (Lorch et al. 2011), and that, *in vivo*, RSC associates most strongly with the first three genic nucleosomes (Yen et al. 2012). Remarkably, our data indicate that much of the information required for this preference must be carried by the selected nucleosomes themselves, without the need for a more complex promoter structure, including nearby transcription factor-binding sites. Moreover, the choice made by RSC among isolated mono-nucleosomes *in vitro* is indeed highly relevant to transcription in living cells, as it preferentially recognizes and remodels mono-nucleosomes that originate from the genes it regulates *in vivo*.

We also find that the preference of RSC for these nucleosomes correlates with the presence of H2A.Z, and that H2A.Z is indeed required to help establish this preference. A role for H2A.Z in stimulating chromatin remodeling has been shown for the ISWI family of chromatin remodelers (Goldman et al. 2010), but it remains to be investigated how H2A.Z facilitates RSC action, and whether it does so by recruitment, catalytic stimulation, or by alternative means. We note that previous experiments on chromatin assembled *in vitro* showed that a mono-nucleosome containing H2A.Z was less readily remodeled than a canonical nucleosome, not only by RSC, but also by other chromatin remodelers tested (Li et al. 2005). One possible explanation for the discrepancy between the outcome of these previous experiment and our results is that the reconstitution experiments by Li *et al.* were performed with DNA containing a single, strong nucleosome positioning sequence, which might affect chromatin remodeling in a manner distinct from that used in our study, which used either natural mono-nucleosomes, or a closed circular plasmid containing recombinant nucleosome arrays.

Somewhat against our expectation, we failed to observe a significant correlation of RSC activity with a specific histone modification, such as acetylation. This is in contrast to prior evidence, which showed that acetylation *does* impact RSC function (see, for example, (Chatterjee et al. 2011; Lorch et al. 2011). Indeed, Bartholomew and co-workers reported that histone H3 tail acetylation enhanced RSC recruitment, and that it also increased nucleosome mobilization and H2A/H2B displacement in a bromodomain-dependent manner (Chatterjee et al. 2011). Interestingly, however, while histone acetylation stimulated recruitment of RSC and nucleosome remodeling via both octamer sliding and hexasome formation, it did not markedly stimulate histone eviction/ejection. Thus, rather than contradicting previous findings, our failure to detect a significant effect of histone acetylation might simply be down to the specific histone eviction assay chosen for our study. It is, however, also worth pointing out that having eight bromodomains might provide RSC with remarkable flexibility in detecting and using many different acetylation marks/positions. Therefore, one would not necessarily expect a large reliance on a *single* mark, but possibly rather a general preference for highly modified nucleosomes. Indeed, such nucleosomes are generally found in promoters and around the TSS, which are also the regions preferred by RSC in our study.

Intrinsic nucleosome stability is typically studied with nucleosomes assembled *in vitro* that lack the natural repertoire of DNA sequences and epigenetic marks. As such, it has remained an open question how nucleosomes assembled *in vivo* differ in their intrinsic biochemical stability. Here, we report evidence that some promoters of protein-coding genes and some tRNA coding regions harbor exceptionally unstable nucleosomes. The finding that many tRNA genes contain an intrinsically highly unstable nucleosome is particularly interesting, also in light of the finding that although nucleosomes *were* previously mapped on such genes, these positions seem to be generally under-occupied *in vivo* (Brogaard et al. 2012). Nucleosomes in these positions may well be further depleted during our purification, but those that *are* isolated clearly shed their bound DNA much more readily than other nucleosomes when they are incubated at 30°C for 15-20 minutes, underscoring that they are indeed intrinsically unstable. It seems an obvious possibility that high intrinsic nucleosome instability has evolved on transcribed regions of the very short tRNA genes because it is evolutionarily advantageous.

The intrinsic instability of promoter nucleosomes agrees well with our understanding of gene function as well: promoters will benefit from more fluid (unstable) nucleosomes to facilitate binding of regulatory factors and nucleation of the transcription initiation complex. This stands in line with the recent finding that certain promoter nucleosomes in nuclei are particularly sensitive to digestion by MNase (Kubik et al. 2015). Nevertheless, the intrinsically unstable nucleosomes described here differ operationally from nucleosomes called as ‘unstable’ *in vivo* by ChIP-Seq or MNAse-Seq experiments (see, for example (Kubik et al. 2015), which was suggested to reflect regions devoid of nucleosomes (see (Chereji et al. 2017) and references therein). Taken together, our results suggest that some nucleosomes in unstable genomic regions are intrinsically unstable through characteristics that are carried by the nucleosome through purification. This intrinsic instability is not due to the underlying sequence (alone), but correlates weakly with an increase in a variety of histone marks (Pokholok et al. 2005), including H3K9Ac, H4Ac and H3K4me3. It is tempting to speculate that intrinsic nucleosome stability is determined by the *combination* of the histone modifications and underlying DNA sequence that characterize them rather by any individual feature in isolation.

In conclusion, the experimental system presented here provides a new tool for studying chromatin *in vitro* in a manner that preserves the natural sequences and epigenetic marks. Its chief advantages are that (1) the entire genome can be interrogated simultaneously, (2) the ‘indirect’ influences from processes occurring on chromatin *in vivo* have been removed, and (3) nucleosomes harbor most, if not all, the epigenetic features as they are found inside the cell. While the experimental system was established based on a yeast nucleosome library, and using RSC as an example of the ‘nucleosome selectivity factor’, it should be possible to similarly apply it to nucleosomes and chromatin-associated factors from other cell types.

## MATERIALS AND METHODS

### Purification of yeast genomic chromatin

Nucleosomes were prepared from strain W303 (wild type) or isogenic *htz1*Δ. Nuclei were prepared largely as described in (Almer and Horz 1986). Nucleosomes were prepared from nuclei by digestion with MNase (New England Biolabs); these were subjected to DEAE chromatography and then loaded on a staggered 20-45% sucrose gradient. Fractions containing the final, purified mono-nucleosomes were pooled.

### RSC-dependent nucleosome disassembly assay

The assay was adapted from the protocol described in (Lorch et al. 2006). RSC:Nucleosome molar ratio was 1:4-1:2. After analysis, gel slices were excised, and DNA extracted using a commercial kit (Life Technologies GeneJET). This DNA was used for sequencing or qPCR analysis. The nucleosome array ejection assay is described in (Clapier et al. 2016); the RSC:nucleosome molar ratio was 1:2 and incubation was for 90 min.

### High-throughput sequencing

Adapters were ligated to mono-nucleosomal DNA using the TruSeq ChIPSeq Sample Prep (Illumina), and sequenced on Illumina HiSeq2500.

### Nucleosome analysis

Relative recoveries of individual sequences in the four bands (NUC, DNA, NUCR and DNAR) were determined by standard qPCR using the DNA obtained from a nucleosome disassembly assay. Native chromatin immunoprecipitation was performed using Protein A Dynabeads (Life Technologies); DNA was purified using a commercial PCR purification kit (Life Technologies GeneJET). Antibodies used were all from Abcam: #8580 (H3K4me3), #9050 (H3K36me3) and #1791 (H3). Primers for *YEF3* and *SSP120* were designed according to (Kim and Buratowski 2009).

### Protein purification

TAP-tagged RSC was purified from Rsc2-TAP cells according to (Lorch and Kornberg 2004). His-tagged Nap1 was purified from *Escherichia coli* according to (Hizume et al. 2013).

### Bioinformatic analysis

Complementary paired-end reads were merged using FLASH (Magoc and Salzberg, 2011) and then as single end reads aligned to the sacCer3 genome using Bowtie2 (Langmead and Salzberg 2012). Read counts per sliding window were normalized for GC content with the R package EDASeq (Risso et al. 2011). Windows with fewer than 5 reads in DNA+NUC or DNAR+NUCR in both replicates were excluded. Normalized log-fold changes in the two replicates between the read counts in DNA, and NUC (DNAR and NUCR) per window were computed with the R package DESeq2 (Love et al. 2014). All gene start sites (TSS’s) were taken from Ensembl release 91 (Zerbino et al. 2018). GC-contents were determined from the nucleosome sequence in the reference genome. Poly(dAdT) tracts were defined as at least 5 consecutive A’s or T’s in a sequence and computed in a similar manner. Histone mark data, H2A.Z scores and hotness were obtained from Pokholok et al. (2005), Albert et al. (2007) and Dion et al. (2007), respectively. Nucleosome occupancy data were obtained from Kaplan et al (2009) and Brogaard et al (2012). Gene expression data (RNA-Seq) was obtained from Nagalakshmi et al. (2008). We used HybMap expression data from Parnell et al (2015) to define RSC-target genes.

Details can be found in Supplementary Materials.

## Acknowledgements

This work was supported by the Francis Crick Institute (which receives its core funding from Cancer Research UK (FC001166, FC010110), the UK Medical Research Council (FC001166, FC010110), and the Wellcome Trust (FC001166, FC010110)) and by a grant from the European Research Council, Agreement 693327 (TRANSDAM). N.M.L. is a Winton Group Leader in recognition of the Winton Charitable Foundation’s support towards the establishment of the Francis Crick Institute. NML is additionally funded by a Wellcome Trust Joint Investigator Award (103760/Z/14/Z), the MRC eMedLab Medical Bioinformatics Infrastructure Award (MR/L016311/1) and core funding from the Okinawa Institute of Science & Technology Graduate University. A.H.E. was supported by an EMBO long-term fellowship. We thank Dr. Tim Parnell for his kind assistance with processed datasets; Drs. Barbara Davis/Roger Kornberg (Stanford University) and Maggie Kasten (Cairns lab) for the kind gift of RSC protein; Dr. Christoph Kurat (Diffley Lab, FCI) for providing Nap1; The FCI Advanced Sequencing Facility for high-throughput sequencing; Dr. Judith Zaugg (EMBL) for sharing datasets; and Drs. Borbala Mifsud, Anna Poetsch and Sebastian Steinhauser (Luscombe lab) for valuable advice on the computational analysis.

## Authors contributions

Idea and conceptualization: AHE and JQS. AHE performed all experiments (Svejstrup lab), except the nucleosome array ejection assays, which were performed by CRC (Cairns lab). All computational analysis was performed by AC (Luscombe lab), with contributions from AHE and ED. JQS, NML and BRC supervised different aspects of the work. AHE, AC, and JQS wrote the paper, with input from all authors.

## Supplemental Figures and Legends

**Figure S1.**
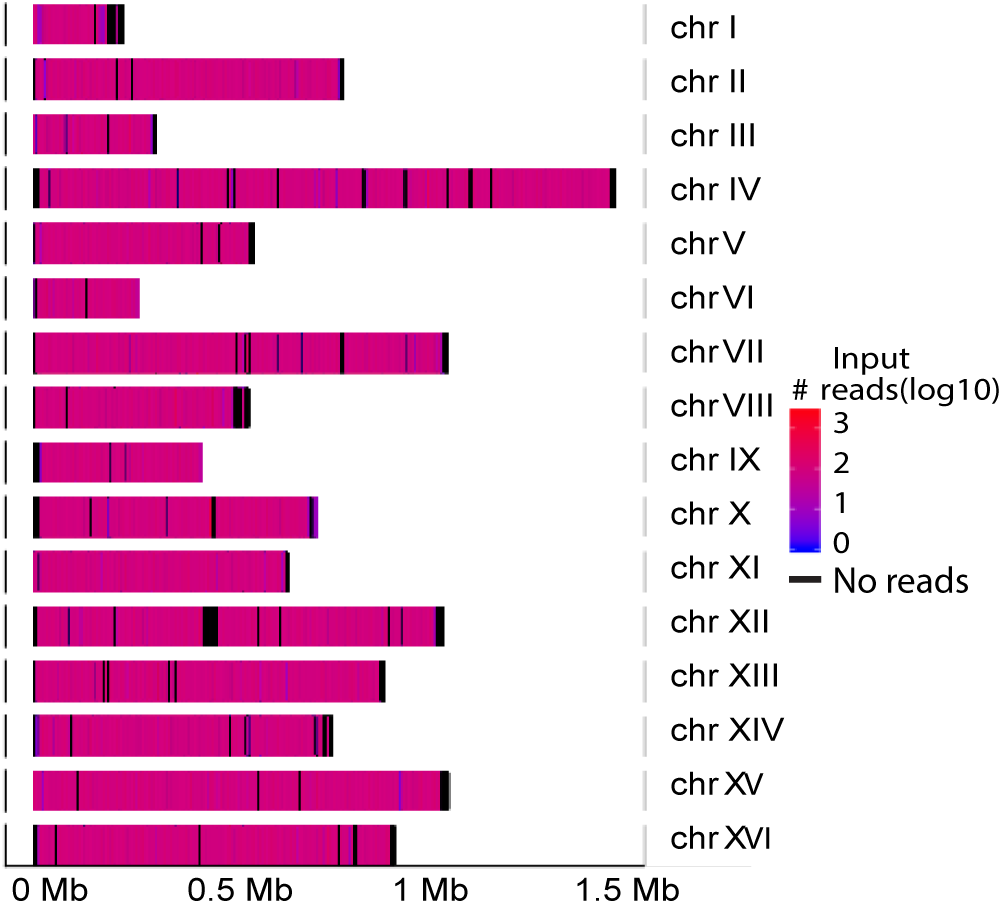
Most of the genome is represented in the mono-nucleosome library. Read density in the input material across the different yeast chromosomes is shown.

**Figure S2.**
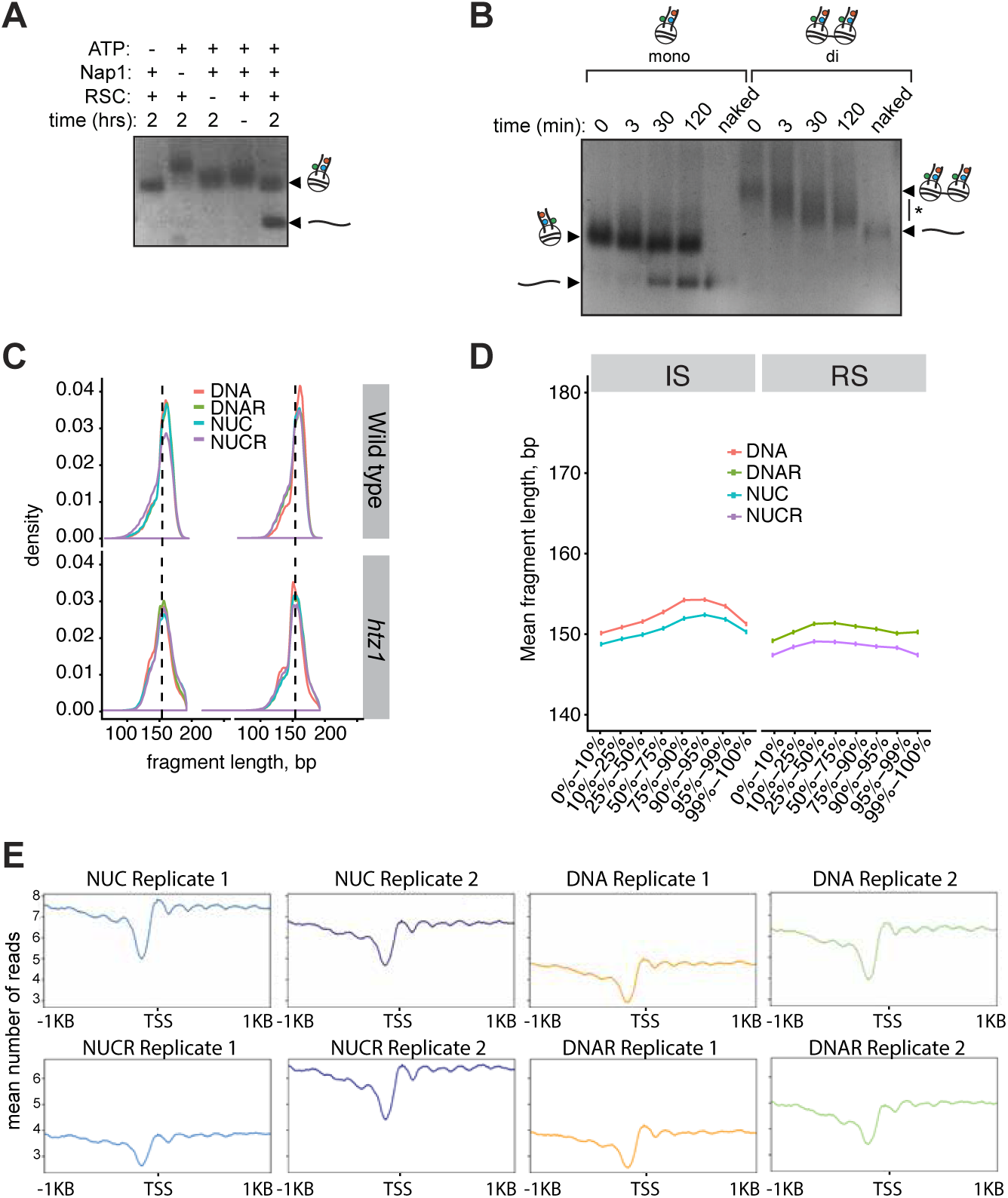
Characterization of assay and nucleosomes. **A.** Nucleosome disassembly depends on ATP, Nap1 and RSC. The assay was performed as for Figure 2B and bands separated by agarose gel-electrophoresis. **B.** Reaction time-course of RSC-dependent nucleosome disassembly of mono- and di-nucleosomes. Nap1 and ATP were included in excess. “Naked” shows naked DNA. Asterisk shows smear of hybrid species consisting of mono-nucleosomes and rearranged di-nucleosomes. **C.** DNA fragment lengths after trimming, mapping and removal of PCR duplicates in the different categories, in nucleosome libraries from WT and *htz1*, respectively. Labels are as shown in Figure 2A.**D.** Average DNA fragment lengths as in **C** in the different IS and RS categories, respectively. IS and RS categories were assigned using the maximal overlapping window for each read. **E.** Average profile of mapped reads around the TSS in the different replicates is highly similar, and resembles that observed by, for example, (Kaplan et al. 2009).

**Figure S3.**
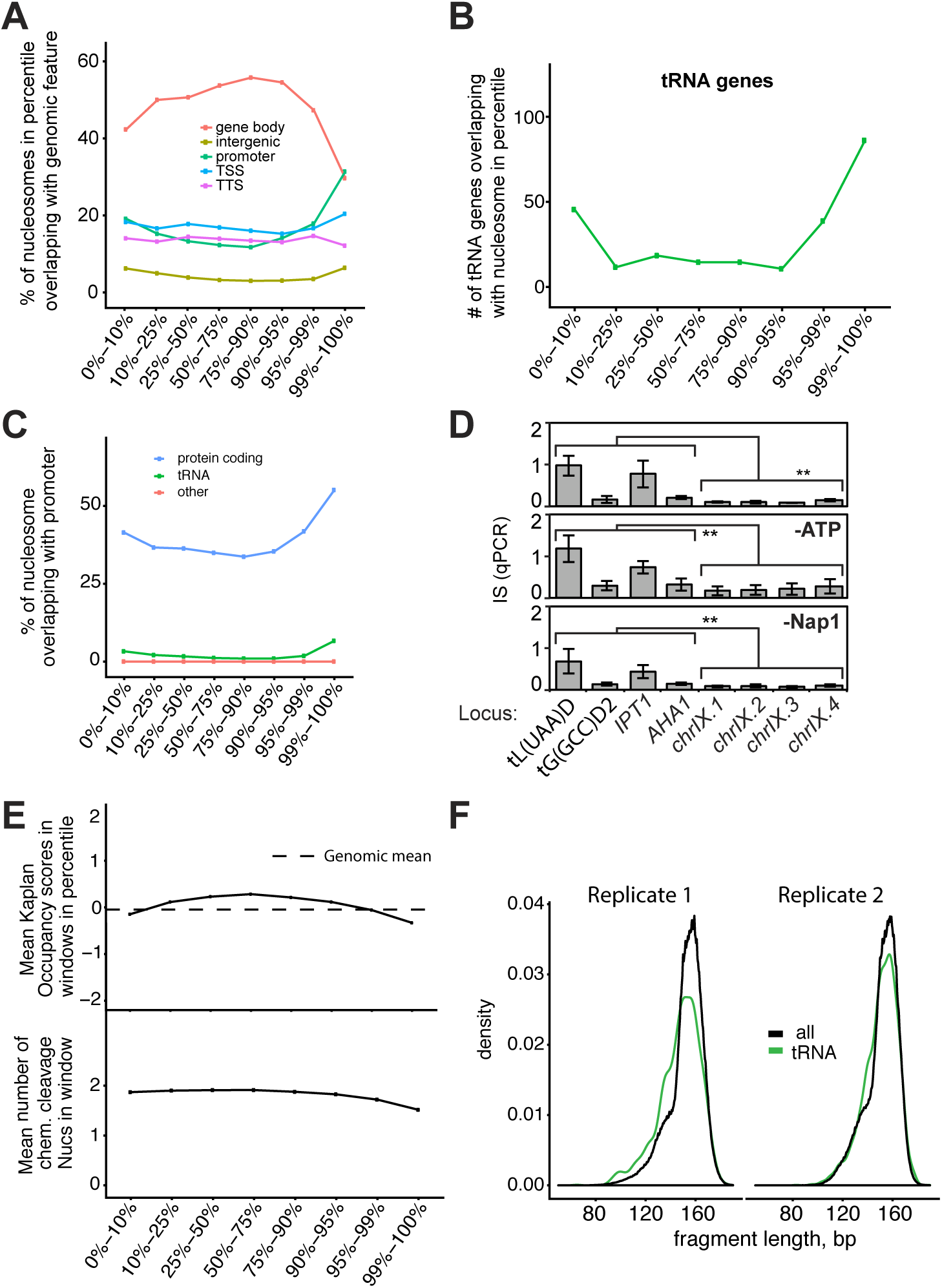
Characterization of intrinsically unstable nucleosomes. **A.** IS scores plotted against nucleosomes that overlap with genomic features, as indicated. **B.** Number of tRNA genes that overlap with a nucleosome found in the different IS percentiles. **C.** IS scores plotted against nucleosomes in the promoter region of genes, as indicated. **D.** Instability as measured in this assay does not require ATP or Nap1. Analysis as in Figure 2G. Asterisks show statistical significance by Wilcoxon t-test (**= p < 0.01). **E.** Number of nucleosomes detected by (Kaplan et al. 2009)(upper) and (Brogaard et al. 2012)(lower) across the different IS percentiles. **F.** DNA fragment lengths at tRNA genes compared to all other fragments.

**Figure S4.**
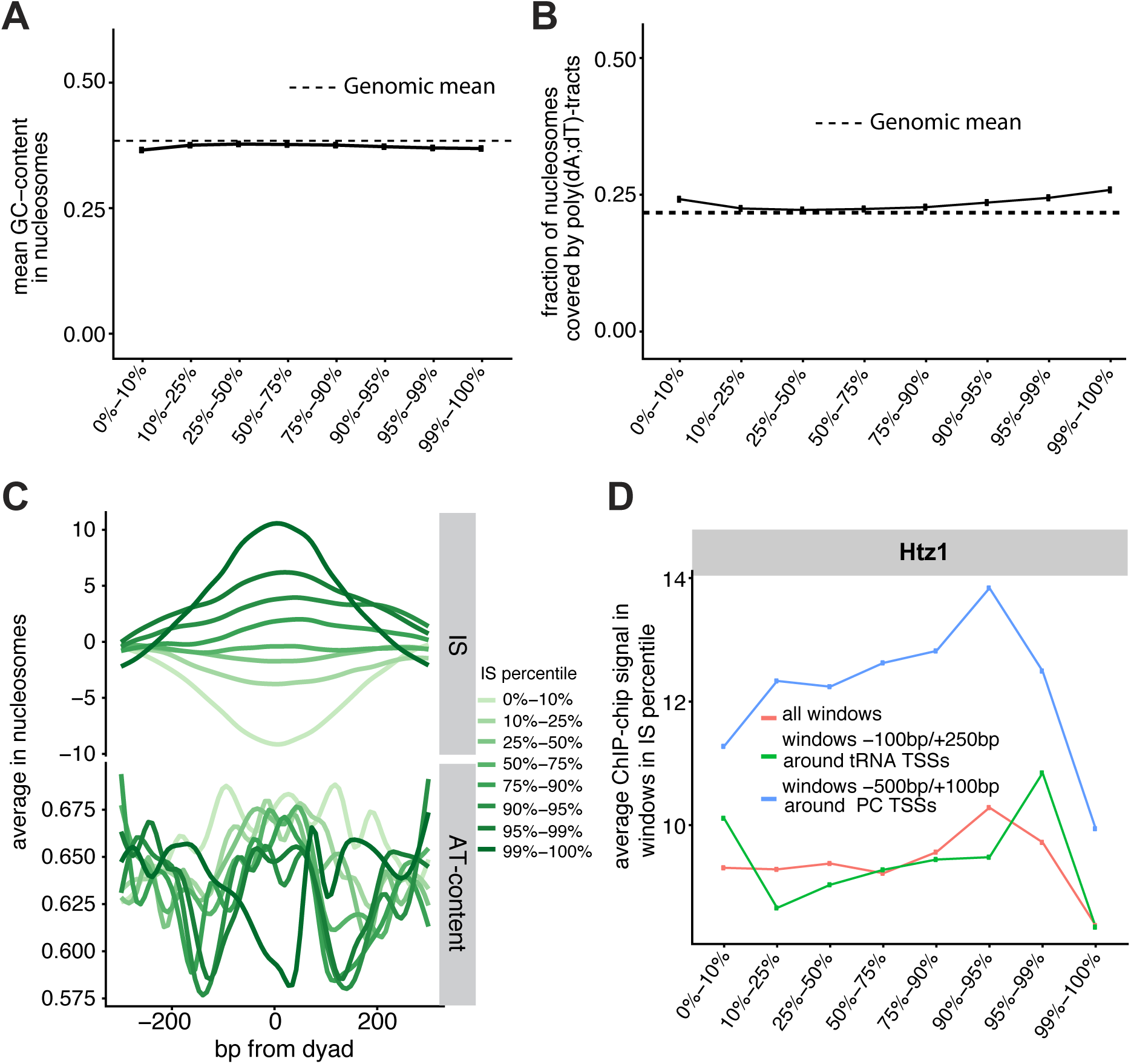
Features characteristics of intrinsically unstable nucleosomes. **A.** Mean GC content in nucleosomes across the different IS percentiles compared to that of the genome average. **B.** Average percent of nucleosomes covered by poly(dA:dT) tracts across the different IS percentiles compared to that of the genome average. **C.** Upper, IS scores across the different percentiles, centered on the dyad of the nucleosome (indicated by 0). Lower, poly(dA:dT) tracts across the same. **D.** H2A.Z density according to (Albert et al. 2007) across the different IS percentiles, at genes or overall.

**Figure S5.**
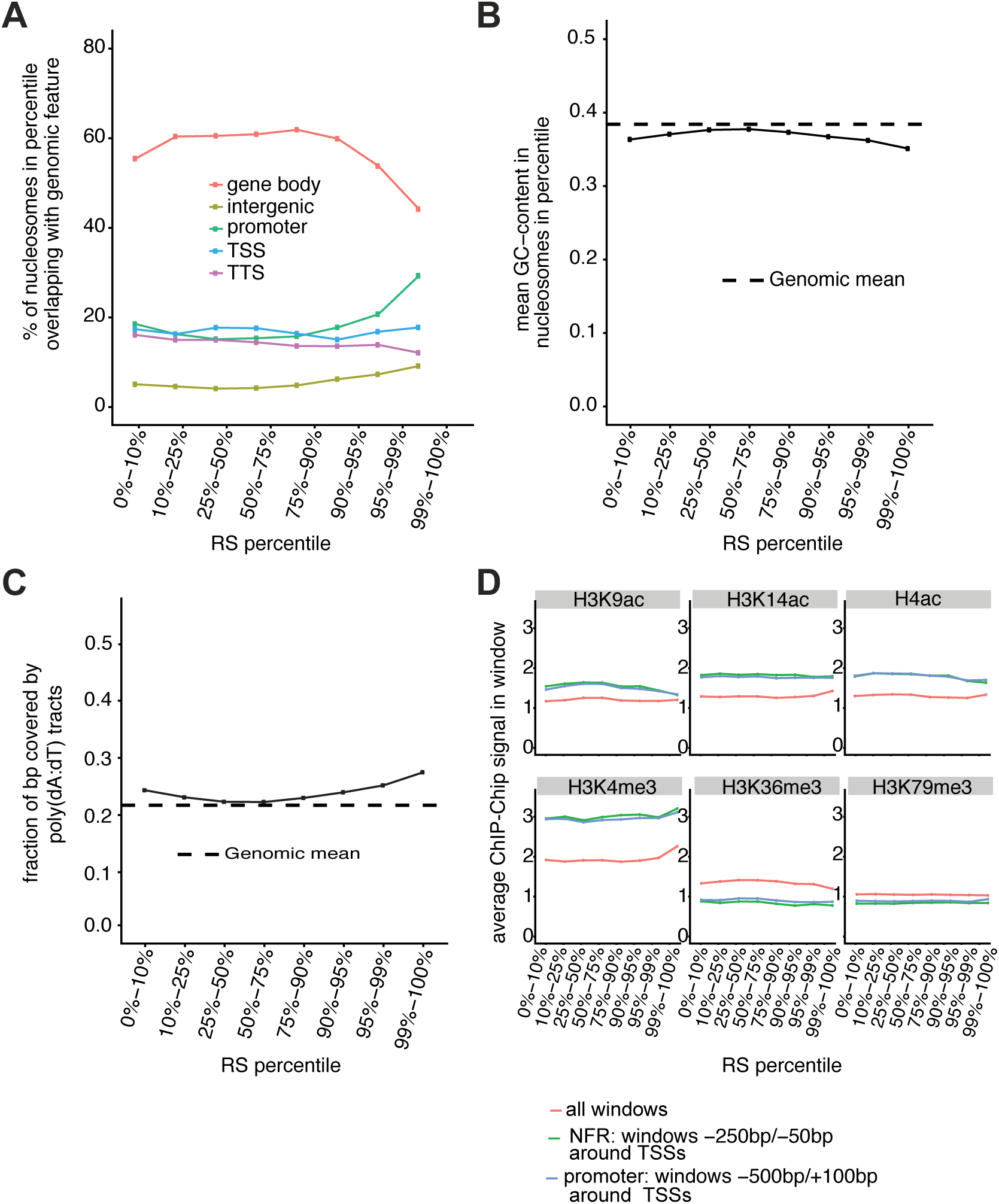
Features characteristics of nucleosomes remodeled by RSC. **A.** IS percentiles plotted against nucleosomes that overlap with genomic features, as indicated. **B.** Mean GC content across the different RS percentiles compared to that of the genome average. **C.** Average percent of nucleosomes covered by poly(dA:dT) tracts across the different RS percentiles compared to that of the genome average. **D.** Histone marks detected by (Pokholok et al., 2005) at genes and overall, across the different RS percentiles.

**Figure S6.**
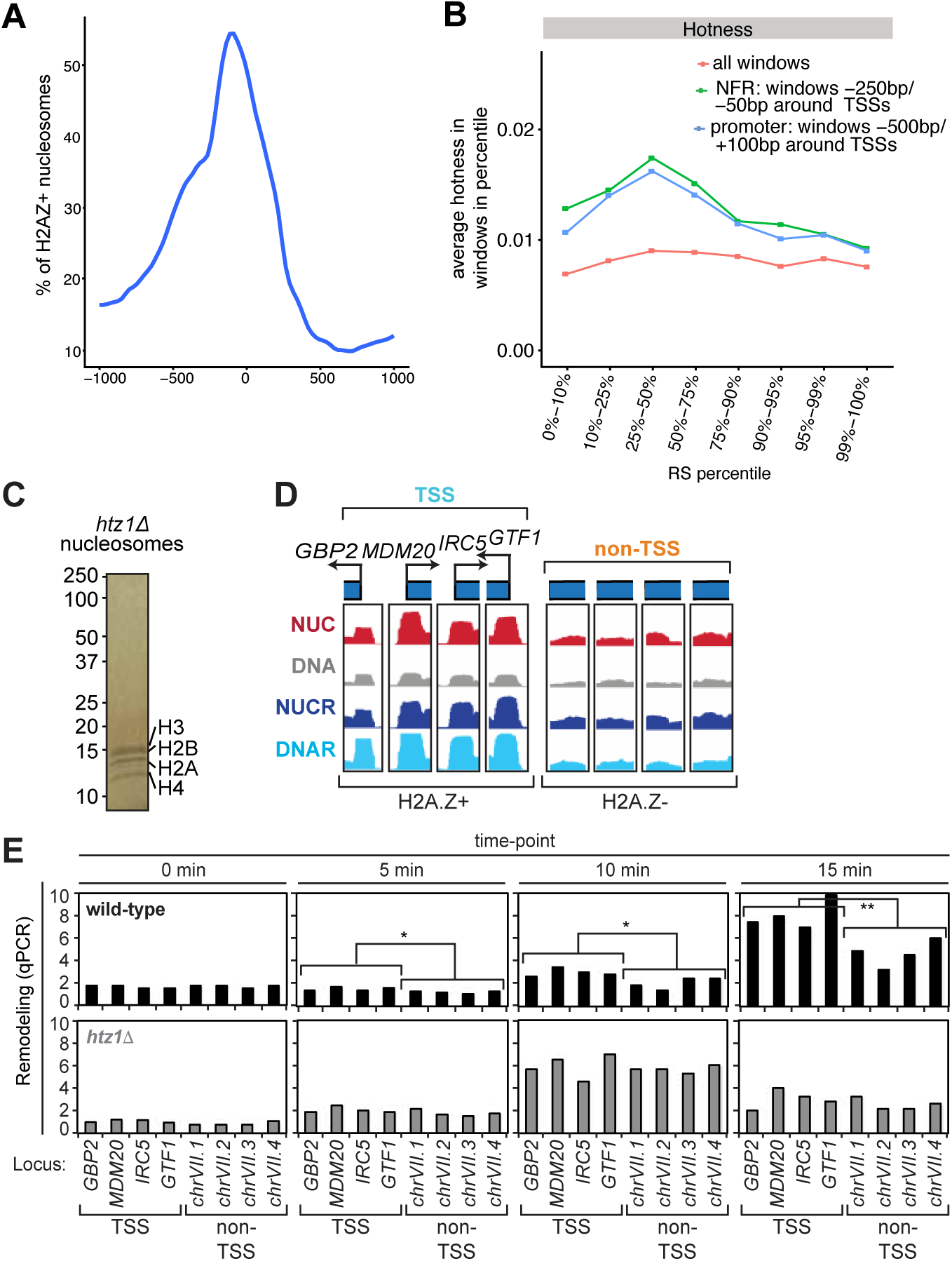
H2A.Z is characteristics of nucleosomes remodeled by RSC. **A.** H2A.Z-containing nucleosomes and their position around the TSS of genes, using data from (Albert et al. 2007). **B.** Nucleosome hotness across the different RS percentiles, at NFRs and promoters compared to that of the genome average, using data from (Dion et al. 2007). **C.** Nucleosomes used for qPCR analysis. **D.** Nucleosomes used for q-PCR experiments on the effect on RSC remodeling of containing high (H2A.Z+) or low (H2A.Z-) H2A.Z density. **E.** Data for Figure 5C decomposed into individual nucleosomes. Note that for the experiments in Figure 5C, nucleosomes that are not on the TSS were used as controls, given that (1) the background activity of RSC on all nucleosomes is already high, and (2) many TSS’s that we call as lacking H2A.Z might actually carry enough H2A.Z to stimulate catalysis, since nearly all genes have been reported to carry some H2A.Z on their promoters (Raisner et al. 2005; Albert et al. 2007).

**Figure S7.**
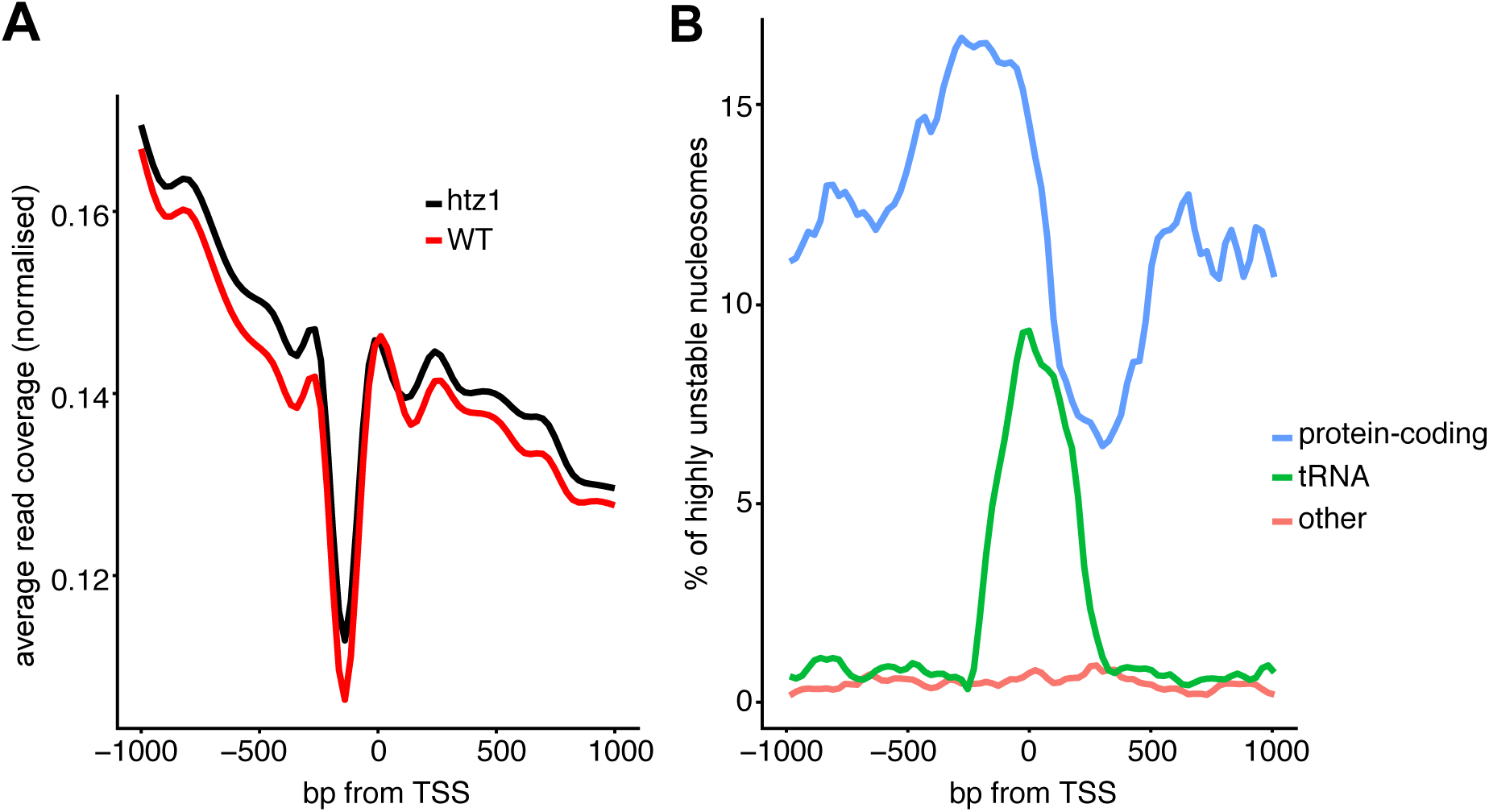
Characteristics of nucleosome libraries from *htz1Δ* cells. **A.** Average read coverage comparison between repeat libraries from WT and *htz1Δ*. **B.** Profile of nucleosome occupancy in the 99th IS percentile, around the TSS of different gene types. Note the overall similarity to wild-type (Figure 3C).

## SUPPLEMENTARY MATERIALS AND METHODS

### Purification of yeast genomic chromatin

Nuclei were prepared by an adaptation of the protocol described in (Almer and Horz 1986). Six litres of YPD were inoculated with 300 ml saturated starter culture in the morning to harvest cells grown to mid-log phase (4-6 cells/ml) in the early afternoon. Cells were pelleted at 3,500 x g for 10 minutes and resuspended in 240 ml (50 mM Tris pH 7.5, 40 mM DTT) for gentle shaking at 30°C for 15 minutes. Cells were pelleted at 11,000 x g for 5 minutes (JA-10 rotor) and resuspended in 60 ml 0.5x YPD/1 M sorbitol at room temperature (from here on all buffers were supplemented with standard protease inhibitors and 5 mM sodium butyrate and 5 mM trichostatin A as deacetylase inhibitors). For spheroblasting, 100 mg of zymolase 20T were added, and the cells shaken slowly for 30-50 minutes at 30°C, until the OD_600_ of a 1:100 dilution in 1% SDS was less than 5% of the initial OD_600_ (e.g. 0.034 vs. initial 0.330). It is important not to overdigest, since the histone tails are sensitive to proteolysis. Zymolysis was stopped by addition of 100 ml cold 1 M sorbitol, and all subsequent operations were performed in the cold. Spheroblasts were pelleted and rinsed twice by centrifuging at 1,500 x g for 5 minutes (JA-10 rotor) and washing twice with 200 ml cold 1 M sorbitol. The rinsed spheroblasts were resuspended in 100 ml Ficoll Buffer (18% Ficoll 400, 20 mM potassium phosphate pH 6.8, 1 mM MgCl_2_, 0.25 mM EDTA, 0.25 mM EGTA) to lyze cells and release the nuclei. Nuclei were centrifuged at 45,000 x g for 20 minutes (Ti-45 rotor), then resuspended in 30 ml SucB1 (0.34 M sucrose, 20 mM Tris pH 7.5, 50 mM KCl, 5 mM MgCl_2_). The resuspended nuclei were then layered on 15 ml SucB2 (1.7 M sucrose, 20 mM Tris pH 7.5, 50 mM KCl, 5 mM MgCl_2_) and centrifuged at 115,000 x g for 30 minutes (SW-32 rotor) to further clean up the nuclei. These were resuspended in 10 ml SucB1, aliquoted to ∼12 x 1 ml, and centrifuged at 18,000 x g for 1 minute. Supernatants were discarded, and pelleted nuclei frozen in liquid nitrogen for storage. Frozen nuclei could be stored indefinitely at −80°C.

Nucleosomes were prepared from nuclei over two days. On the morning of day 1, one aliquot of nuclei was thawed and resuspended in 500 µl cold MNase Buffer (50 mM NaCl, 13 mM Tris pH 8.0, 6.4 mM CaCl_2_, 0.2 mM EDTA, 0.2 mM EGTA), before adding 10 µl 0.1 M CaCl_2_ and protease and deacetylase inhibitors as above. Resuspended nuclei were pre-heated for 90 seconds at 37°C, and digestion performed with ∼40,000 U MNase (∼20 µl of 2,000 U/µl from New England Biolabs) for 2 minutes at 37°C. Digestion was stopped by addition of 25 µl 0.5 M EGTA and placing of the tube on ice for 2 minutes. Optimal MNase concentrations were determined in a trial digestion for each batch of nuclei, by testing five different concentrations of MNase with 100 µl resuspended nuclei and choosing the concentration that yields mostly mono-nucleosomes, but still results in formation of some oligo-nucleosomes. It is crucial that the digestion be performed as fast as possible, and that all other steps be performed on ice or in the cold room to avoid proteolysis of histone tails. After digestion, the incubated nuclei were spiked with more protease inhibitors, then centrifuged at 18,000 x g for 1 minute. The supernatant containing mono-nucleosomes was kept and centrifuged one more time to remove remaining nuclei. The solution should be brown to yellow in colour, and clear. For DEAE chromatography, we used a self-poured 1.2 cm x 7 cm column (6 ml bed volume) equilibrated in DEAE-NucB-20 (20 mM NaCl, 10 mM Tris pH 7.5, 1.5 mM MgCl_2_, 10% glycerol). All 500 µl of nuclear lysate were loaded at 0.3 ml/min, and elution was performed in a 20−800 mM NaCl gradient in DEAE-NucB at 0.2 ml/min over 40 minutes. Chromatin eluted at about 400 mM in the second half of the major peak (See Figure 1B). The high salt concentration of ∼400 mM used for elution from DEAE ensured that DNA-binding transcription factors were released and washed off, while the gradient centrifugation served to remove contaminants and separate free nucleosomes from nucleosomes bound by other proteins. Before proceeding with the sucrose gradient, fractions containing chromatin were identified with a quick agarose gel analysis by extracting DNA from 20 µl using the Qiagen PCR purification kit, performing a brief digestion with RNAse A and separating bands on a 1.5% agarose gel. Fractions containing chromatin were pooled and loaded on a staggered 20-45% sucrose gradient consisting of 5 ml each of 45, 40, 35, 30, 25 and 20% sucrose in Sucrose Buffer (30 mM NaCl, 10 mM Tris pH 7.5, 1 mM EDTA, 1 mM EGTA). Ultracentrifugation was performed at 200,000 x g for 26 hours (SW41 rotor). Fractions of ∼500 µl were then collected by piercing the bottom of the tube (∼8 drops per fraction, ∼20 fractions total). To identify the fractions containing DNA, 15 µl of each fraction was incubated with ∼30 µg RNAse A for 20 minutes at 37°C, before adding 3 µl loading buffer containing 1% SDS, and separating bands on a 1.5% agarose gel. Fractions containing mono-nucleosomes were pooled, distributed into 25 or 50 µl aliquots, flash-frozen in liquid nitrogen, and stored at −80°C. Final recoveries were ∼1 to 1.5 ml at concentrations of ∼20 ng/µl, corresponding to 20-30 µg of DNA. Each aliquot was thawed only once. We recommend testing the integrity of the histone tails by performing a western blot for histone H3 and checking for the absence of a lower band corresponding to proteolyzed histone. DNA concentrations in the final material were determined by qPCR using a standard curve of two different PCR products of known concentration that covered a part of the genome (primers P764/765 and P788/789).

### Strains used in this study

Nucleosomes were prepared from strains W303 (wild-type) and *htz1Δ* from the Saccharomyces Genome Deletion Project (Brachmann et al. 1998).

### RSC-dependent nucleosome disassembly assay

The assay was adapted from the protocol described in (Lorch et al. 2006), except that DNA was stained with ethidium rather than by radioactive labelling. Each reaction was performed in 20 µl volumes, with 6-10 µl nucleosomes, 800 ng RSC, 2 µg Nap1, 1 mM ATP, 20 mM potassium acetate pH 7.6, 15 mM Hepes pH 7.9, 3 mM MgCl_2_, 75 µg/ml bovine serum albumin, and protease inhibitors (Rsc:nucleosome molar ratio ∼1:4 – 1:2). After incubation for 20 minutes at 30°C, 1 µl of ∼2 mg/ml plasmid DNA was added to capture free RSC, followed by the addition of 5 µl 30% glycerol. At this point, the mixture could be flash-frozen in liquid nitrogen and stored indefinitely at −80°C. For native gel-electrophoresis, the thawed mixture was loaded on a pre-cooled 2% agarose gel in 0.5x TBE, then run for ∼30 minutes at 50 V and ∼1.5 hours at 100 V. Bands were stained by soaking the gel in 1 µg/ml ethidium bromide for one hour, destained with water for 30 minutes and bands analyzed with UV light. In our experience, bands were generally very weak, with their intensity varying greatly depending on the concentration of nucleosomes. After analysis, gel slices were excised, and DNA extracted using a commercial kit (Life Technologies GeneJET), to be eluted into 50 µl elution buffer. This DNA was suitable for sequencing or qPCR analysis.

### High-throughput sequencing

Adapters were ligated to mono-nucleosomal DNA using the TruSeq ChIPSeq Sample Prep (Illumina), amplified in 12 cycles, and sequenced in 4- or 5-plex multiplex format on Illumina HiSeq 2,500. The number of reads obtained were: 50,531,688 (NUC), 40,528,367 (DNA), 68,912,998 (NUCR), 49,153,637 (DNAR) and 54,797,627 (input).

### qPCR analysis of unstable and remodelled nucleosomes

Relative recoveries of individual sequences in the four bands (NUC, DNA, NUCR and DNAR) were determined by standard qPCR using the DNA obtained from a nucleosome disassembly assay. Nucleosomes to investigate for analysis were chosen by scanning the coverage of all four bands for nucleosomes with the desired properties (as in Figure 2E and 2F). IS(qPCR) and RS(qPCR) are defined as IS(seq) and RS(seq), except that the log2 is not taken. Thus, IS(qPCR) = [DNA]/[NUC] and RS(qPCR) = [DNAR]/[NUCR] – IS, where the numbers in brackets are the relative concentrations of a given sequence as determined by qPCR.

### Native chromatin immunoprecipitation

Protein A Dynabeads (25 µl, Life Technologies) were washed with PBS, pre-loaded with 4 µl antibody, then washed again twice with PBS. After resuspension in 300 µl IP-B2 (150 mM NaCl, 50 mM Tris pH 8.0, 2 mM EDTA, 0.1% NP-40 and 0.01% SDS), 50 µl nucleosomes were added. 50 µl of the input were kept for qPCR analysis, and the remaining 300 µl nutated for 3 hours at 4°C. Nucleosome-bound beads were washed three times with 500 µl IP-B2 and DNA eluted by incubating with 50 µl Elution-B (IP-B2 with 1% SDS) for 20 minutes at room-temperature. 100 µl TE pH 8.0 were added to the eluates and to the input, and the DNA purified using a commercial PCR purification kit (Life Technologies GeneJET). Sequence quantitation by was performed by standard analysis by quantitative PCR. Antibodies used were all from Abcam: #8580 (H3K4me3), #9050 (H3K36me3) and #1791 (H3). Primers for *YEF3* and *SSP120* were designed according to (Kim and Buratowski 2009).

### Read processing and alignment

The quality of reads was checked with FASTQC (available online at https://www.bioinformatics.babraham.ac.uk/projects/fastqc/). Reads were trimmed with trimgalore (parameters: `--length 0`) (available online at www.bioinformatics.babraham.ac.uk/projects/trim_galore/). Complementary paired-end reads were combined into a single sequence covering the entire nucleosome using FLASH (parameters: `-m 10 -M 53`) (Magoc and Salzberg 2011) and then as single end reads aligned to the sacCer3 genome using Bowtie2 (parameters: `default`)(version 2.3.3.1, (Langmead and Salzberg 2012)). The resulting SAM file was converted into BAM format using Samtools (Li et al. 2009). Duplicate reads were identified using Picard (parameters: `default`) (available online at http://broadinstitute.github.io/picard/) and subsequently removed using Samtools. Only fragments that mapped uniquely with a mapping quality of at least 30 and that were longer than 50 bp and shorter than a nucleosome with a long linker (190 bp) were retained. Reads mapping to the mitochondrial chromosome were discarded. The final numbers of fragments for each band in each replicate (15 min and 20 min incubation) after trimming are shown in the table below.

|  | **Wild-type**<br># fragments<br>Replicate 1/ Replicate 2 | **Htz1**<br># fragments<br>Replicate 1/ Replicate 2 |
| --- | --- | --- |
| **DNA** | 465,006 / 607,212 | 202,079 / 588,507 |
| **NUC** | 729,397 / 688,374 | 279,007 / 687,732 |
| **DNAR** | 377,768 / 505,395 | 360,910 / 948,949 |
| **NUCR** | 408,719 / 657,549 | 426,977 / 750,861 |

### Sliding windows, Instability and Remodelling Scores and nucleosome regions

We excluded both the first and last 1000 bp of each chromosome of the yeast genome. We divided the yeast genome into sliding windows of size 167bp with step size 25bp and allocated any overlapping mapped reads to the windows. As previously reported for MNase-Seq for nucleosomal DNA (Chung et al. 2010), we observed an enrichment for GC-rich reads, and we therefore normalized the read counts for the GC content of that window with the R package EDASeq (Risso et al. 2011). We excluded windows with fewer than 5 reads in DNA+NUC or DNAR+NUCR in both replicates from further analysis. Read counts per window were subsequently normalized for sequencing depth and normalized log-fold changes in the two replicates between the read counts in DNA and NUC (DNAR and NUCR) per window were computed with the R package DESeq2 (Love et al. 2014). After normalization and quality filtering, 450,257 windows were recovered for comparison.

The Instability Score (IS) was defined as the log-fold change between the read counts DNA and NUC for each individual nucleosome, ie. IS = log_2_(DNA/NUC). The Remodeling Score (RS) was defined as RS = log_2_(DNAR/NUCR)-IS. We divided the sets in 8 percentiles (0-10%, 10-25%, 25-50%, 50-75%, 75-90%, 90-95%, 95-99%, 99-100%). Finally, overlapping or adjacent regions in the same percentile for IS or RS were merged to obtain unstable and remodeled regions. The median widths of these regions (median width ∼ 240 for IS, ∼220 for RS) are too small to contain more than one nucleosome, and we refer to these regions as nucleosomes in the remainder of the text for simplicity. To compare fragment lengths in the 4 bands (Figure S2D), we used the maximal overlapping window to assigned each read an IS/RS percentile.

### Comparison with published datasets

After annotating the unique nucleosome map of the Widom lab (Brogaard et al. 2012) to the yeast genome sacCer3 using the R package rtracklayer (Lawrence et al. 2009), nucleosomes were defined as 83 bp on either side of the dyad, i.e. the nucleosomes span 167 bp to accommodate average sized linker for sacCer. Nucleosome occupancy data from the Segal lab (Kaplan et al. 2009) was annotated to the sacCer3 version of the yeast genome using the R package rtracklayer (Lawrence et al. 2009).

### Enrichment of genetic and epigenetic features

We used the *biomaRt* package (Durinck et al. 2005; Durinck et al. 2009) to retrieve gene annotation information from Ensembl release 91 (Zerbino et al. 2018) using BioMart web services (Kasprzyk et al. 2004; Smedley et al. 2015). GC-contents were determined from the nucleosome sequence in the reference genome sacCer3 (The Bioconductor (The Bioconductor Dev Team 2014). Poly(dAdT) tracts were defined as at least 5 consecutive A’s or T’s in a sequence and computed in a similar manner. Histone mark data were obtained from (Pokholok et al. 2005). They were assigned to each window or nucleosome by first smoothing them using a rolling window of size 500 with step size 100 across the genome and then assigning their average value over a given window or nucleosome as score. H2A.Z scores and hotness were obtained from (Albert et al. 2007) and (Dion et al. 2007), respectively, then assigned to each nucleosome/window in a similar manner.

### Metaprofiles and Heatmaps

To produce the metaprofiles of the raw reads, we used deepTools (Ramirez et al. 2016) to remove the GC bias of the raw read counts (paramters: `default`), and subsequently to center reads and convert the resulting bam file into a BigWig file (parameters: `-- minFragmentLength 140 –maxFragmentLength 190 –centerReads –binSize 1`), and then to plot the metaprofiles. Other metaprofiles were built by using the regions of interest (TSSs or nucleosomes) as templates in which the presence/absence of a feature (window, nucleosome, poly(dA:dT) tracks) is noted by 1/0. When the nucleosomes were analysed in a gene context (promoter or TSS), they were first flipped accordingly to the sense of transcription. When plotting metaprofiles of the IS or RS scores, these binary vectors were multiplied with the IS/RS-score of the window. The resulting vectors were then summed and normalised by the total number of features. The metaprofile plots were generates with the R package ggplot2 (Wickham 2016) and smoothed with a polynomial regression with a polynomial function of degree 51. The heatmaps of reads in genomic contexts were generated with the R package ggbio (Yin et al. 2012).

### Genomic features assessment

In order to look at the genomic distribution of the nucleosome regions, we divided them into TSSs, TTSs, promoters (−600 to −100 upstream of TSS according to the sense of transcription), gene bodies and intergenic regions. The promoters were trimmed if they were overlapping a gene body upstream. We first assigned TSS (11,536 IS and 8,582 RS) and TTS (11,389 IS and 8,5,23 RS) or both (2,417 IS and 1,410 RS) nucleosomes; the remaining nucleosomes were intersected with promoters (11,689 IS and 9,702 RS) and then gene bodies (42,433, IS and 35,696, RS). 3,199 IS and 2,715 RS nucleosomes fall in intergenic regions.

### Gene expression analysis

Gene expression data was obtained from (Nagalakshmi et al. 2008) and annotated to the sacCer3 genome using the R package rtracklayer (Lawrence et al. 2009). Further, we used the genomic locations of the HybMap expression data from (Parnell et al. 2015) to compute the log-fold change of expression between the *rsc2-V457M* strain and *wild-type* for those genes in the dataset where at least half of the gene body was covered by probes. We then defined the individual expression values for each gene as the average of the (sense) probes that overlapped with the gene body. We defined genes to be RSC-targets if the absolute value of log-fold change in expression between *rsc2-V457M* strain and *wild-type* was in the 80th percentile of the absolute log-fold changes.

We assigned each of the genes a mean RS as the mean of the scores of the windows intersecting 300bp up- and 50bp downstream of the TSS of each gene respectively. This resulted in 615 genes having both a finite log-fold change and a defined RS score: 557 protein-coding, 9 tRNAs, 43 snoRNAs, 4 ncRNAs, and 2 snRNAs; of which 124 genes are RSC-targets (83 protein-coding, 31 snoRNA, 6 tRNAs, 2 snRNAs and 2 ncRNAs). We then mapped these RS values to the genome wide percentiles to define RS percentiles.

### Nucleosome Ejection Assay

Nucleosome arrays used in the ejection assays were assembled as 100 µl reaction containing 5 µg of pBluescript plasmid DNA mixed with 23 pmol of recombinant histone octamers, containing either canonical H2A or H2A.Z (Htz1) variant, in 2 M KCl, 1x NEB4, 0.1 mg/ml BSA, in the presence of 10 U of Topoisomerase I (NEB) with a linear salt-gradient dialysis applied from 2 M to 50 mM KCl at 30°C, using an Econo-Pump (Bio-Rad) and Slide-A-Lyser Mini Dialysis units with a 7,000 molecular weight cutoff (Thermo Scientific).

Nucleosome ejection assays were performed in a 50 µl reaction by incubating 1 pmol of RSC protein complex (20nM) with 500 ng of nucleosome arrays (described above) in 10 mM Tris buffer, pH 7.4, 50 mM KCl, 3 mM MgCl_2_, 0.1 mg/ml BSA, 1 mM ATP at 30°C with shaking at 500 rpm in a Thermomixer (Eppendorf) for 90 min. Deproteinization was performed by adding 5 µl of proteinase K at 10 mg/ml and 2.5 µl of 20% SDS and incubated at 50°C for 1h. Samples were subsequently precipitated in ethanol, prior to two-dimensional separation on a 1.3% agarose gel as described in (Clapier et al. 2001). Gels were stained for 15 min in a 1 µg/ml ethidium bromide solution and scanned on a Typhoon Trio (Amersham, GE).

